# A homing suppression gene drive with multiplexed gRNAs maintains high drive conversion efficiency and avoids functional resistance alleles

**DOI:** 10.1101/2021.05.27.446071

**Authors:** Emily Yang, Matthew Metzloff, Anna M. Langmüller, Xuejiao Xu, Andrew G. Clark, Philipp W. Messer, Jackson Champer

**Affiliations:** Department of Computational Biology, Cornell University, Ithaca, NY 14853; Department of Molecular Biology and Genetics, Cornell University, Ithaca, NY 14853; Institut für Populationsgenetik, Vetmeduni Vienna, Veterinärplatz 1, 1210 Wien, Austria; Vienna Graduate School of Population Genetics, 1210 Wien, Austria; Center for Bioinformatics, School of Life Sciences, Peking-Tsinghua Center for Life Sciences, Peking University, Beijing, China 100871

## Abstract

Gene drives are engineered alleles that can bias inheritance in their favor, allowing them to spread throughout a population. They could potentially be used to modify or suppress pest populations, such as mosquitoes that spread diseases. CRISPR/Cas9 homing drives, which copy themselves by homology-directed repair in drive/wild-type heterozygotes, are a powerful form of gene drive, but they are vulnerable to resistance alleles that preserve the function of their target gene. Such resistance alleles can prevent successful population suppression. Here, we constructed a homing suppression drive in *Drosophila melanogaster* that utilized multiplexed gRNAs to inhibit the formation of functional resistance alleles in its female fertility target gene. The selected gRNA target sites were close together, preventing reduction in drive conversion efficiency. The construct reached a moderate equilibrium frequency in cage populations without apparent formation of resistance alleles. However, a moderate fitness cost prevented elimination of the cage population, showing the importance of using highly efficient drives in a suppression strategy, even if resistance can be addressed. Nevertheless, our results experimentally demonstrate the viability of the multiplexed gRNAs strategy in homing suppression gene drives.

## INTRODUCTION

At the frontier of pest and disease vector control, gene drives hold the potential to influence large, wild populations. These engineered genetic elements have the ability to spread quickly by biasing inheritance in their favor, allowing for the manipulation of population sizes or traits such as disease transmission^1–5^.

Gene drives can act through many mechanisms and include both engineered and naturally occurring forms^6^. For engineered homing drives, the CRISPR/Cas9 system has been widely used to create gene drive constructs in many organisms, including yeast^7–10^, flies^11–25^, mosquitoes^26–38^, and mice^39^. The homing mechanism converts an organism heterozygous of the drive into a homozygote in the germline, and the drive is thus transmitted to offspring at a rate above 50%. These drives contain a Cas9 endonuclease, which cleaves a target sequence, and at least one guide RNA (gRNA), which directs Cas9 to the cleavage location. The resulting DNA break can be repaired by homology-directed repair (HDR) using the drive allele as a template, thereby copying the drive into the wild-type chromosome.

However, a major obstacle that impedes drive efficiency is the alternative DNA repair method of end-joining, which does not use a homologous template and often alters the target sequence, preventing further recognition by the gRNA/Cas9 system. Such gRNA target site mutations, whether formed by drive cleavage or preexisting in the population, are therefore considered resistance alleles and can form at high rates in the germline as well as in the embryo due to cleavage activity from maternally deposited Cas9 and gRNA^11,17,20–22,24,25,27,31,37,38^. Resistance alleles that disrupt the function of the target gene by causing frameshifts or otherwise sufficiently changing the amino acid sequence tend to be more common in almost all gene drive designs, and we call them “r2” alleles. By contrast, “r1” a **l**eles” preserve gene function and are therefore particularly detrimental to gene drives. If the drive allele imposes a greater fitness cost than the resistance allele, which is usually the case for functional alleles in most drives that target native genes, then the resistance alleles will outcompete the drive and thwart its potential to modify or suppress the population^20,24,26,40–42^.

While modification drives aim to genetically alter a population, for instance by spreading a specific gene variant or genetic cargo, the goal of suppression drives is to ultimately reduce and potentially even eliminate a population, usually by disrupting an essential but haplosufficient gene target and thus impacting a negative fitness impact in drive homozygotes. For example, such a drive could cleave and be copied into a gene with a recessive knockout phenotype that affects viability or fecundity. As the drive increases in frequency in the population (via heterozygotes, which remain fertile and viable^43^), the proportion of sterile or nonviable individuals will increase, thereby reducing population size. Even if the drive forms some nonfunctional resistance alleles, they would show the same phenotype as drive alleles, thus only somewhat slowing the spread of the gene drive and likely still allowing successful suppression^44^. Functional resistance alleles, on the other hand, would be expected to have a drastic effect on this type of drive, quickly halting and reversing population suppression and outcompeting the drive^26,37,45–47^. Therefore, the success of a suppression drive hinges on its ability to reduce the functional allele formation rate to a sufficiently low level while also avoiding gRNA targets where functional alleles are already present in the population.

The formation of such functional resistance alleles was successfully prevented in one *Anopheles* study targeting a highly conserved sequence of a female fertility gene, since end-joining repair of such a target would be unlikely to result in a functional mutation^32^. However, the population size in this experimental study was necessarily limited to several hundred individuals^28,32,35^, so it remains unclear if any functional resistance alleles could still form against this drive in much larger and more variable natural populations. Additional measures may thus be needed in a large-scale release to prevent the formation of functional resistance alleles. Furthermore, such highly conserved sequences in possible target genes for suppression drives may not be available in other species, and even high conservation of the target site alone is sometimes insufficient to prevent formation of functional resistance alleles, as shown by another recent study in *Anopheles*^30^.

Multiplexing gRNAs has been proposed as a mechanism that could reduce the rate of functional allele formation by recruiting Cas9 to cleave at multiple sites within the target gene. If one gRNA target is repaired by end-joining in a way that leaves the gene functional, additional sites could still be cleaved, resulting in additional opportunities for drive conversion or creation of nonfunctional mutations. Simultaneous cleavage at multiple sites and repair by end-joining could also result in large deletions, which would usually render the target gene nonfunctional^20^. Several models indicate that multiplexed gRNAs would likely be effective at reducing functional resistance alleles^24,48,49^, and a handful of experimental studies have supported this notion^11,20,24,25^. Furthermore, multiplexing of gRNAs is capable of increasing drive conversion efficiency, as has been demonstrated in a modification homing drive with two gRNAs^20^. However, one study using four gRNAs for a homing suppression drive reported very low drive efficiency^11^, which would likely prevent effective population suppression^24,47^, particularly in larger, spatially-structured populations^45,50^. This reduction in efficiency was in part caused by repetitive elements in the drive, which resulted in removal of large portions of the drive by recombination during homology directed repair^11^. However, widely spaced gRNAs also likely played an important role, since failure to cleave the outermost gRNAs would require end resection of large DNA tracts before an area of homology would be reached with the drive allele^24^.

These findings suggest that an effective suppression drive could consist of multiple gRNAs targeting closely spaced sequences. The best target would likely be a female-specific haplosufficient but essential fertility gene^45^. Although such a drive would impose a high fitness cost to homozygous females, it could still spread at a high rate through germline conversion in heterozygous females and males, and any nonfunctional resistance alleles would eventually be removed from the population rather than outcompeting the drive. Females with any combination of drive and nonfunctional resistance alleles would be infertile.

Here, we construct such a drive in *Drosophila melanogaster* with four multiplexed gRNAs targeting *yellow-g*. The homing suppression drive demonstrated in these experiments showed elevated inheritance rates and successfully persisted in cage populations that averaged over 4,000 flies per generation without apparent formation of functional resistance alleles. However, the drive also imposed an unintended fitness cost of unknown type. This, together with the low drive conversion rate and high embryo resistance allele formation rate compared to *Anopheles* drives, ultimately prevented suppression of the experimental populations.

## METHODS

### Plasmid construction

The starting plasmids TTTgRNAtRNAi^24^, TTTgRNAt^24^, and BHDcN1^20^ were constructed previously. For plasmid cloning, reagents for restriction digest, PCR, and Gibson assembly were obtained from New England Biolabs; oligos and gBlocks from Integrated DNA Technologies; 5-α competent *Escherichia coli* from New England Biolabs; and the ZymoPure Midiprep kit from Zymo Research. Plasmid construction was confirmed by Sanger sequencing. A list of DNA fragments, plasmids, primers, and restriction enzymes used for cloning of each construct can be found in the Supplemental Information section. We provide annotated sequences of the final drive insertion plasmid and target gene genomic region in ApE format at github.com/MesserLab/HomingSuppressionDrive (for the free ApE reader, see biologylabs.utah.edu/jorgensen/wayned/ape).

### Generation of transgenic lines

Embryo injections were provided by Rainbow Transgenic Flies. The donor plasmid HSDygU4 was injected into *w*^*1118*^ flies along with plasmid TTTygU4, providing the gRNAs for transformation, and pBS-Hsp70-Cas9 (140 ng/µL, from Melissa Harrison & Kate O’Connor-Giles & Jill Wildonger, Addgene plasmid #45945) as the source of Cas9 for transformation. Flies were housed with BDSC standard cornmeal medium in a 25°C incubator on a 14/10-hour day/night cycle.

### Genotypes and phenotypes

Flies were anesthetized with CO_2_ and screened for fluorescence using the NIGHTSEA adapter SFA-GR for DsRed and SFA-RB-GO for EGFP. Fluorescent proteins were driven by the 3xP3 promoter for expression and easy visualization in the white eyes of *w*^*1118*^ flies. DsRed was used as a marker to indicate the presence of the split drive allele, and EGFP was used to indicate the presence of the supporting *nanos*-Cas9 allele^22^.

### Cage study

For the cage study, flies were housed in 30×30×30 cm (Bugdorm, BD43030D) enclosures. The ancestral founder line that was heterozygous for the split drive allele and homozygous for the supporting *nanos*-Cas9 allele was generated by crossing successful transformants with the Cas9 line^22^ for several generations, selecting flies with brighter green fluorescence (which were likely to be Cas9 homozygotes) and eventually confirming that the line was homozygous for Cas9 via PCR.

These flies (heterozygous for the split drive and homozygous for Cas9), together with *nanos*-Cas9^22^ homozygotes of the same age, were separately allowed to lay eggs in eight food bottles for a single day. Bottles were then placed in cages, and eleven days later, they were replaced in the cage with fresh food. Bottles were removed from the cages the following day, the flies were frozen for later phenotyping, and the egg-containing bottles returned to the cage. This 12-day cycle was repeated for each generation.

### Artificial selection small cage study

For the small cage population experiment designed to detection functional resistance alleles, flies heterozygous for the split drive and homozygous for Cas9 were crossed to each other. We then crossed three batches of 50 drive heterozygous males to 50 drive heterozygous females, which were allowed to lay eggs for two days. Their progeny were collected three times each day at six-hour intervals after they started eclosing. Non-fluorescent flies (indicating absence of the drive allele) were discarded. Over 90% of females were clearly virgins by visual phenotype using this collection scheme. Thirteen days after the first egg laying day, the original vial was discarded, and fourteen days afterward, the progeny were allowed to lay eggs for two days after being split randomly in two separate vials. The cycle was then repeated for each generation. With this method, wild-type alleles are removed from the population at an increased rate each generation, compensating for a drive’s intermediate drive conversion rate and fitness cost. This increases the genetic load (suppressive power) of the drive, raising the chance that the population is eliminated instead of reaching an equilibrium frequency, as would be predicted if drive conversion is low (particularly in the presence of high fitness costs and embryo resistance allele formation). If functional resistance alleles form, however, they would usually have high viability, preventing suppression. Thus, lack of population elimination in this experiment would likely indicate that functional resistance alleles were present.

### Phenotype data analysis

Data were pooled into two groups of crosses (drive heterozygous females with *w*^*1118*^ males and drive heterozygous males with *w*^*1118*^ females) in order to calculate drive inheritance, drive conversion, and embryo resistance. However, this pooling approach does not take potential batch effects (offspring were raised in different “batches” - vials with different parents) into account, which could bias rate and error estimates. To account for such batch effects, we conducted an alternate analysis as in previous studies^24,25,51^. Briefly, we fit a generalized linear mixed-effects model with a binomial distribution (by maximum likelihood, Adaptive Gauss-Hermite Quadrature, nAGQ = 25). This model allows for variance between batches, usually resulting in slightly different parameter estimates and increased standard error estimates. Offspring from a single vial were considered as a separate batch. This analysis was performed with the R statistical computing environment (3.6.1) including packages lme4 (1.1-21, https://cran.r-project.org/web/packages/lme4/index.html) and emmeans (1.4.2, https://cran.r-project.org/web/packages/emmeans/index.html). The script is available on Github (https://github.com/MesserLab/Binomial-Analysis). The resulting rate estimates and errors were similar to the pooled analysis (Data Sets S1-3).

### Genotyping

For genotyping, flies were frozen, and DNA was extracted by grinding single flies in 30 µL of 10 mM Tris-HCl pH 8, 1mM EDTA, 25 mM NaCl, and 200 µg/mL recombinant proteinase K (ThermoScientific), followed by incubation at 37°C for 30 minutes and then 95°C for 5 minutes. The DNA was used as a template for PCR using Q5 Hot Start DNA Polymerase from New England Biolabs with the manufacturer’s protocol. The region of interest containing gRNA target sites was amplified using DNA oligo primers YGLeft_S_F and YGRight_S_R. This would allow amplification of wild-type sequences and sequences with resistance alleles but would not amplify full drive alleles with a 30 second PCR extension time. After DNA fragments were isolated by gel electrophoresis, sequences were obtained by Sanger sequencing and analyzed with ApE software (http://biologylabs.utah.edu/jorgensen/wayned/ape).

Deep sequencing and analysis was performed by Azenta Life Sciences on a pool of approximately 100 newly eclosed flies after DNA purification and amplification with the same primers as described above. PCR products were treated with enzymes for 5’ Phosphorylation and dA-tailing, and T-A ligation was performed to add adaptors, and products were ligated to beads. PCR was conducted using primers on the adaptors, and the final library was purified and qualified by beads. Qualified libraries were pair-end sequenced for 150 nucleotides using Illumina Hiseq Xten/Miseq/Novaseq/MGI2000. Data was analyzed with cutadapt (1.9.1), flash (v1.2.11), bwa (0.7.12-r1039), and Samtools (1.6).

### Fitness cost inference framework

To quantify drive fitness costs, we modified a previously developed maximum likelihood inference framework^25,52^. Similar to a previous study^53^, we extended the model to two unlinked loci (drive site and a site representing undesired mutations from off-target cleavage that impose a fitness cost). The Maximum Likelihood inference framework is implemented in R (v. 4.0.3)^54^ and is available on GitHub (https://github.com/MesserLab/HomingSuppressionDrive).

In this model, we make the simplifying assumption of a single genetic loci and a single gRNA at the gene drive allele site. Each female randomly selects a mate. The number of offspring generated per female can be reduced in certain genotypes if they have a fecundity fitness cost, and the chance of a male being selected as a mate can be reduced if they have a mating success fitness cost. In the germline, wild-type alleles in drive/wild-type heterozygotes can potentially be converted to either drive or resistance alleles, which are then inherited by offspring. At this stage, wild-type alleles at the off-target site are also cleaved, becoming disrupted alleles that may impose a fitness cost. The genotypes of offspring can be adjusted if they have a drive-carrying mother. If they have any wild-type alleles, then these are converted to resistance alleles at the embryo stage with a probability equal to the embryo resistance allele formation rate. This final genotype is used to determine if the offspring survives based on viability fitness.

We set the germline drive conversion rate and the embryo resistance allele formation rate to the experimental inferred estimates (76.7% for drive conversion using the average of male and female rates and embryo cut rate of 52.2%, see results section - note that we did not include in this average the data from females in the drier vials as described in the results section since their progeny had lower viability, which would make assessment of drive conversion unreliable if based only on drive inheritance). Based on previous observations^19,20,22,24^, we set the germline nonfunctional formation rate to 22.2 % so that nearly all wild-type alleles would either be converted to a drive allele or a resistance allele. Functional resistance alleles were not initially modeled since they are expected to be extremely rare in the 4-gRNA design (but see below). Note that in this framework, drive conversion and germline resistance allele formation take place at the same temporal stage in the germline. We set the germline cut rate at the off-target locus to 1 and did not model additional off-target cuts in embryos with drive-carrying mothers. This represents the simplest model of mostly distant off-target sites that are mostly cut in the germline when Cas9 cleavage rates are highest (actual off-target cleavage would likely be at many sites at much lower rates, with some linked to the drive alleles, which would not be possible to easily model with our maximum likelihood method^53^). We assumed that in drive carriers at the beginning of the experiment, 50% of the off-target sites are cut because the drive carrier flies all came from male drive heterozygotes. All drive carriers were initially drive heterozygotes. In future generations, we used the relative rate of drive heterozygotes and homozygotes (among drive carriers with DsRed) as well as relative rates of other genotypes with a wild-type (non-DsRed) phenotype as predicted in the maximum likelihood model.

In one model, we assumed the fitness costs would occur only in female drive/wild-type heterozygotes due to somatic Cas9 expression and cleavage. In the remaining scenarios, we assumed that drive fitness costs would either reduce viability or reduce female fecundity (separately from the sterility of female drive homozygotes) and male mating success. These fitness costs either stemmed directly from the presence of the drive or from cleavage at a single off-target site (representing multiple possible off-target sites that were unlinked to the drive). Our fitness parameters represent the fitness of drive homozygotes (or simply the net fitness of drive heterozygotes for the somatic Cas9 cleavage fitness model). Heterozygous individuals were assigned a fitness equal to the square root of homozygotes, assuming multiplicative fitness costs between loci and alleles. The model incorporates the sterility of females not carrying any wild-type allele of *yellow-g*, and thus, any inferred fitness parameters < 1 represent additional fitness costs of the drive system.

To estimate the rate at which resistance alleles might be functional types, we took the best model for each cage and introduced a new “relative r1 rate” parameter, representing the fraction of resistance alleles that become functional alleles instead of nonfunctional alleles. Our germline rate of 22.2% then became the total resistance allele formation rate, while the experimental measured embryo nonfunctional resistance allele formation rate remained fixed at 52.2%. This relative r1 rate parameter was then inferred as above to obtain an estimate and confidence interval.

## RESULTS

### Drive construct design

In this study, we aim to develop a population suppression homing drive in *D. melanogaster* that utilizes multiple gRNAs to improve drive efficiency and reduce the rate of functional resistance allele formation. Our drive construct targets *yellow-g*, which has previously been used as a female fertility homing suppression drive target in flies^11^ and mosquitoes^27^. Located on chromosome 3, it is highly conserved across *Drosophila* species (Table S1). The *yellow* gene family is closely related to the major royal jelly family in *Apis mellifera* and has been shown to play a critical role in the membrane proteins of the embryo during egg development in *Drosophila*^55^. Null mutations of *yellow-g* usually result in sterile females when homozygous but show no effects on males or on females when one wild-type copy is present (Figure 1). Both integration of the drive or formation of nonfunctional resistance alleles that disrupt gene function will result in such null alleles. Conversion of wild-type alleles to drive alleles in the germline of drive heterozygotes allows the drive to increase in frequency in the population (Figure 1). This will lead to an increasing number of sterile individuals that can eventually induce population suppression.

**Figure 1.**
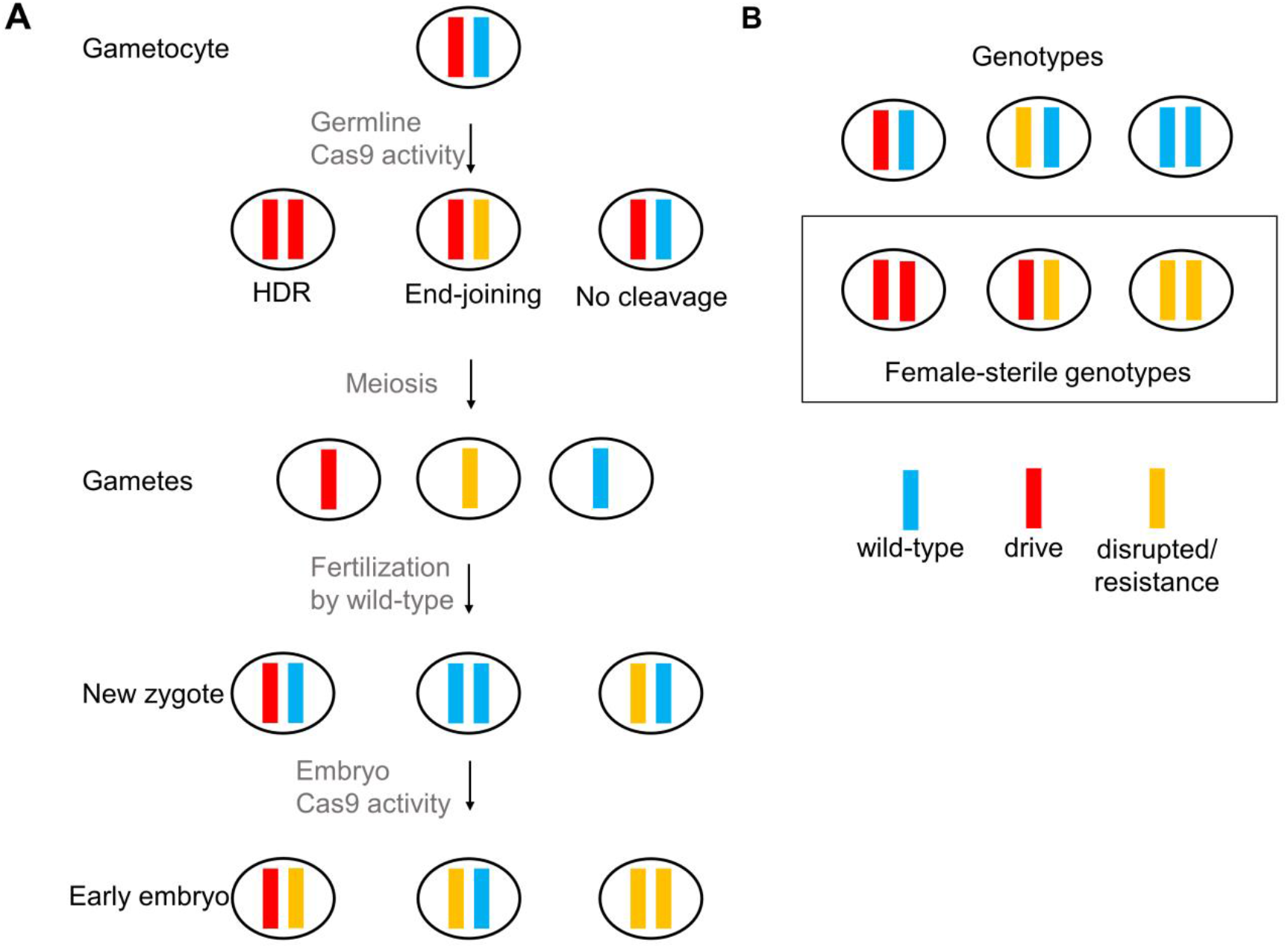
Homing suppression drive inheritance. (**A**) Germline Cas9 activity can convert wild-type allele to drive alleles, though end-joining repair can produce resistance alleles. Maternally deposited Cas9 and gRNA can form additional resistance alleles in the early embryo. (**B**) Females carrying only drive alleles or nonfunctional resistance alleles are sterile.

The drive is inserted between the leftmost and rightmost gRNA target sites of *yellow-g*, providing the template for homology directed repair (Figure 2). The drive construct contains a DsRed fluorescent marker driven by the 3xP3 promoter for expression in the eyes to indicate the presence of a drive allele. It also contains four gRNAs (confirmed to be active by target sequencing) within tRNA scaffolding that target the second exon of *yellow-g*. By eliminating the need for multiple gRNA cassettes, the construct is more compact and avoids repetitive gRNA promoter elements. All four gRNA target sites are located within the second exon of *yellow-g*, a site chosen to allow for closely spaced target sites in a moderately conserved area (Table S1) while still being sufficiently far from the end of the gene to ensure that frameshift mutations would always disrupt the gene’s function. This design should both increase the drive’s homing rate as well as the probability that when a resistance allele is formed, it is an nonfunctional allele (disrupting the target gene’s function) rather than an functional allele. gRNA sites were also chosen to avoid strong off-target sites (Table S2). The Cas9 element, required for drive activity, is placed on chromosome 2R and provided through a separate line that carries Cas9 driven by the *nanos* germline promoter and EGFP with the 3xP3 promoter. In this split-Cas9 system, the drive will only be active in individuals where the Cas9 allele is also present^22^.

**Figure 2.**
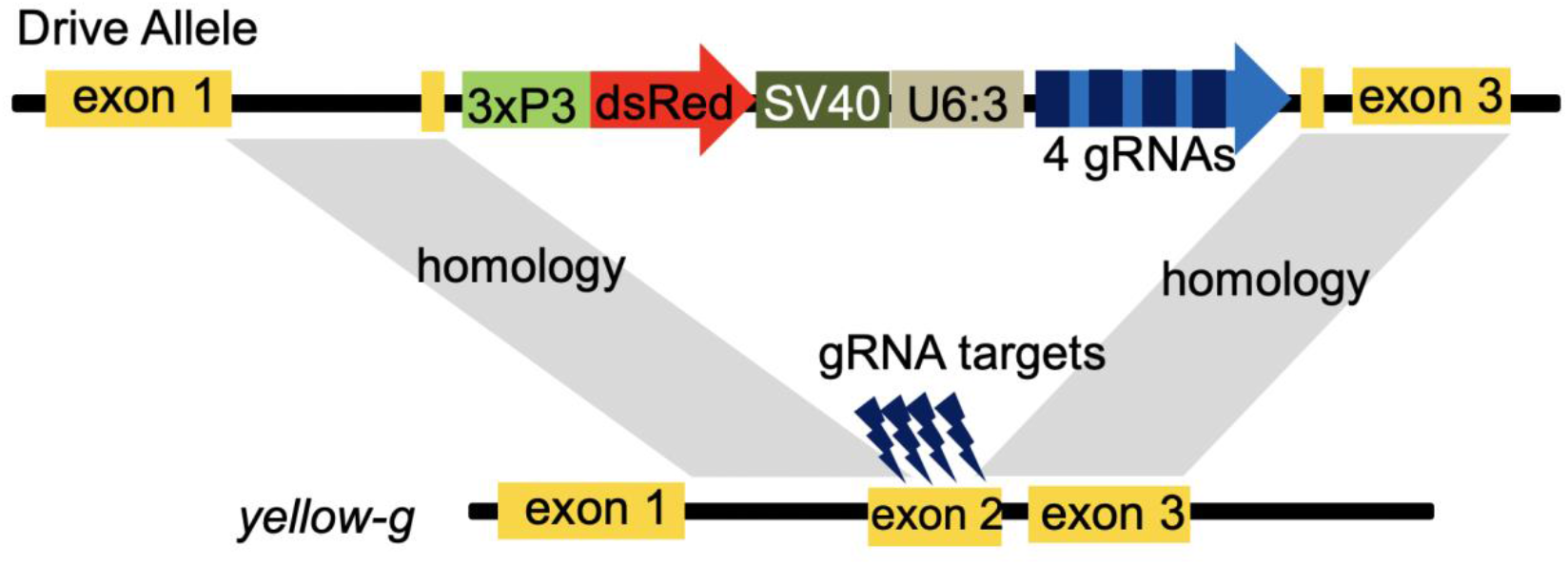
Homing suppression drive schematic. The drive is placed inside the *yellow-g* gene at the gRNA target sites to allow for homology directed repair. A DsRed fluorescence marker is driven by the 3xP3 promoter. Four gRNAs (multiplexed in tRNA scaffolding and driven by the U6:3 promoter) target regions of the second exon of *yellow-g*. This is a split drive system, so Cas9 (driven by the *nanos* promoter) was provided at an unlinked site in the genome for drive experiments.

### Drive inheritance

Successful transformants were used to establish fly lines with the construct, which were maintained by removing wild-type females in each generation. We first crossed the drive line to a line that was homozygous for *nanos*-Cas9. The offspring of this cross that had DsRed are expected to carry one copy each of the drive allele and the Cas9 allele, and these flies were then crossed with *w*^*1118*^ flies for drive conversion assessment. The offspring of this second cross were phenotyped for red fluorescence, indicating the presence of the drive allele (Figure 3) and in a subset of the vials, also for green fluorescence, indicating presence of the Cas9 allele. The drive was inherited at a rate of 86.4% in the progeny of female drive heterozygotes (Data Set S1), substantially higher than the Mendelian inheritance rate of 50% (Fisher’s Exact Test, *p* < 0.00001) and thus indicative of strong drive activity. For the progeny of male drive heterozygotes, the inheritance rate was 90.4% (Data Set S2), which was also substantially higher than the Mendelian expectation (Fisher’s Exact Test, *p* < 0.00001). Because we do not expect this drive to reduce the viability of any eggs (except those laid by sterile females) as confirmed for this set of crosses (though see viability section below for an additional data set), we can calculate the rate at which wild-type alleles were converted to drive alleles based on the drive inheritance rate. This drive conversion rate was 72.7% for female heterozygotes and 80.7% for male heterozygotes. These rates were greater than most similar single gRNA designs^19,20,22^ and comparable to similar 2-gRNA designs^20,24,25^, as predicted by a model of multiple gRNAs^24^.

**Figure 3.**
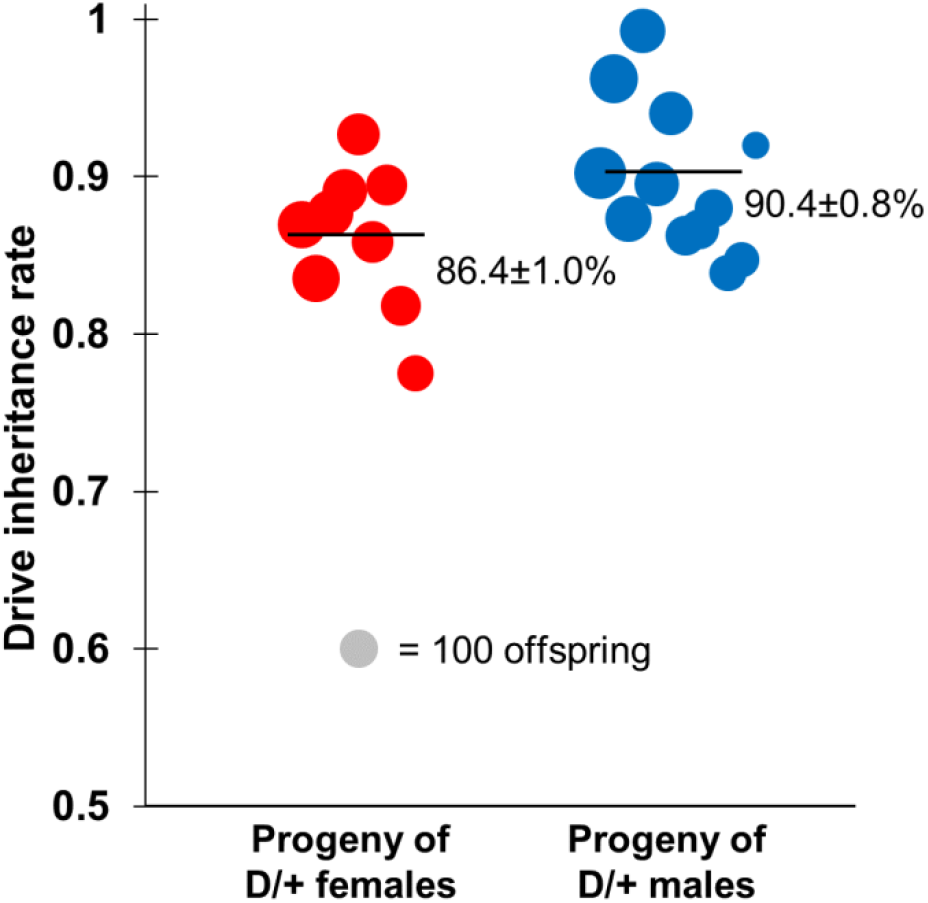
Drive inheritance rates. Drive inheritance as measured by the percentage of offspring with DsRed fluorescence from crosses between drive individuals (heterozygous for the drive and for a Cas9 allele) and wild-type flies. Each dot represents offspring from one drive parent, and the size of dots is proportional to the number of total offspring from the parent. Rate and standard error of the mean are displayed for the overall inheritance rate for all flies pooled together. An alternate analysis that accounts for potential batch effects yielded overall similar rates with slightly increased error estimates (Data Sets S1-2).

The inheritance rate of the Cas9 allele (which should have unbiased inheritance) was 46.7% for females and 43.8% for males. These rates were not significantly different from the Mendelian expectation of 50% (*p* = 0.3 for females and *p* = 0.1 for males, Binomial test), consistent with little to no fitness costs for the Cas9 cassette.

### Resistance alleles and fertility

To determine the rate of resistance allele formation in the embryo due to maternally deposited Cas9 and gRNAs, DsRed female offspring were assessed for fertility. These individuals were daughters of drive heterozygous mothers (heterozygous for both drive and Cas9 alleles and crossed to *w*^*1118*^ males as described above) and could thus have developed embryo resistance. This would convert these flies from fertile drive/wild-type heterozygotes into drive/nonfunctional resistance allele heterozygotes, which are expected to be sterile. Several of these daughters of drive heterozygous mothers were each crossed to two *w*^*1118*^ males. Their vials were observed one week later and compared to control crosses with *w*^*1118*^ females. Vials with no offspring were considered to have sterile females due to embryo resistance allele formation or other factors.

Twelve out of 22 (54.5%) assessed females were sterile, which is significantly higher than the 5% sterility rate (in one out of 20 individuals tested) of female drive-carrier offspring from male drive and Cas9 heterozygotes crossed to *w*^*1118*^ females (*p* < 0.001, Fisher’s exact test). Assuming that this 5% sterility rate represents a baseline for our laboratory flies under the given experimental conditions, we can calculate that an embryo resistance allele formation rate of 52.2% will account for the increased sterility rate in the progeny of drive females. This should provide an estimate for the rate at which the paternal wild-type alleles of *yellow-g* were cleaved at one or more gRNA target sites in embryos with drive mothers.

To analyze resistance alleles from the perspective of individual sequences, progeny from drive heterozygous females or males were Sanger sequenced around the target site (Table S3). To examine embryo resistance, 27 drive-carrying progeny of female drive heterozygotes were analyzed. These individuals received a wild-type allele from their father, which could then undergo cleavage due to maternally deposited Cas9 and gRNAs. We found that ten of these progeny harbored resistance alleles, but only one with a large deletion did not have any gRNA target sites that remained wild-type. Six progeny were fully wild-type, while another nine were mosaic to various degrees. Only some of these mosaics were likely sterile, so these results are approximate agreement with our estimate of 52% sterile progeny, assuming resistance alleles were all nonfunctional (40% with a full resistance sequence, meaning that about 1/3 of the mosaics are likely sterile to reach 52%). Ten non-drive progeny of either female or male drive heterozygotes were also assessed, and half of them were found to be carrying resistance sequences, only one of which did not have any wild-type sites available for cleavage (Table S3). Looking at all sequenced individuals, it is clear that the first and last gRNA cut sites experienced relatively high rates of cleavage, while the middle two gRNA sites experienced low rates of cleavage. A similar 4-gRNA drive with a different target site had the highest activity in the first and third gRNA^24^, and the gRNAs in this drive were placed in the same relative order in the gRNA gene (with the outermost target sites as the first two gRNAs in the gene). Thus, the observed variance in gRNA activity in both studies is likely due to differences in individual gRNA activity levels rather than position in the target site or in the gRNA gene.

To better understand resistance allele formation in the germline, several female and male drive heterozygotes were crossed to each other, and approximately 100 progeny were generated in two vials. All of these were pooled and deep sequenced around the gRNA target sites. Each wild-type allele therefore experienced potential germline cleavage in either male or females, and then experienced further cleavage in the embryo due to maternally deposited Cas9. In contrast to our Sanger sequences of alleles that only underwent potential cleavage in the embryo, we saw substantial cut rates at all four gRNA target sites, including the two middle ones that had very low embryo cut rates (Figure S1). This is generally consistent with the notion that germline cut rates are quite high, with most, though not all, wild-type alleles being converted to resistance alleles or undergoing successful drive conversion by homology-directed repair. Examining individual resistance allele sequences, we found many instances of deletions between cut sites (Figure S2), indicative of cleavage at multiple gRNA sites before end-joining repair and subsequent loss of DNA between the cut sites. Such deletions are potentially disadvantageous in terms of reducing future ability to perform drive conversion. However, they have the benefit of further reducing the chance of functional resistance allele formation because larger deletions are more likely to disrupt the function of the target gene, even if they are in frame.

### Fecundity and viability

Drive homozygous females (as confirmed by sequencing) were found to be sterile as expected. One important issue with population suppression gene drives is leaky somatic expression that can convert drive/wild-type heterozygotes partially or completely into drive/resistance allele heterozygotes (or perhaps even drive/drive homozygotes) in somatic cells, which was responsible for substantially reducing the fertility of mosquitoes carrying homing suppression drives in previous studies^26,32,37^. To determine if drive heterozygotes had altered fertility, three-day old female virgins that were heterozygous for the drive and Cas9 alleles were crossed with *w*^*1118*^ males and then allowed to lay eggs for three consecutive days in different vials, with the eggs counted each day. They laid an average ± standard deviation of 33±4 apparently normal eggs per day (Data Set S1), which was significantly higher than the 20±2 eggs per day laid by *w*^*1118*^ females crossed to drive and Cas9 heterozygous males (Data Set S2, *p* = 0.008, t-test) or the 23±2 eggs per day laid by *w*^*1118*^ females crossed with *w*^*1118*^ males (Data Set S3, *p* = 0.017, t-test). This greater number of eggs per day was likely a batch effect from perhaps slightly older or healthier drive females compared to the *w*^*1118*^ females used. Indeed, if the first day of egg-laying is discounted, the new average of 25±3 eggs per day for drive heterozygous females is statistically indistinguishable from the other groups, regardless of whether the first day of egg-laying is retained in these groups (*p* > 0.1 for all comparisons, t-test). This indicates that any drive cleavage from leaky somatic expression is sufficiently low, such that it does not substantially reduce female fertility (though we cannot rule out small reductions). These results are consistent with the notion that the *nanos*-Cas9 allele has little to no leaky somatic expression, as shown in previous *Drosophila* studies^19,20,22,25^. The offspring of these crosses (females heterozygous for the drive and Cas9 crossed with *w*^*1118*^ males, males heterozygous for the drive and Cas9 crossed with *w*^*1118*^ females, and *w*^*1118*^ fly crosses) did not exhibit any apparent developmental fitness costs. In particular, there were no differences in egg or pupae viability between these three groups of offspring (Data Sets S1-S3).

To increase our sample size of individual crosses to better detect small fitness effects, similar crosses were performed with drive heterozygous females and males together with *w*^*1118*^ individuals, as well as control crosses with only *w*^*1118*^ flies (Data Sets S1-S3). However, these crosses took place at a different laboratory with different food preparation technique. While the food was still able to support flies, it was notably drier. Perhaps because of this, the results of these crosses were notably different for female drive heterozygotes (Data Set S1). Specifically, the drive inheritance of adult progeny was lower (79% in the new batch versus 87% in earlier crosses), and the viability of eggs was also lower (77% in the new batch versus 83% earlier). This reduction in egg viability reached statistical significance compared to a new batch of control females bearing only one Cas9 allele that were tested in identical conditions (*p* = 0.0001, Fisher’s Exact Test), which themselves had the same egg viability as earlier *w*^*1118*^ controls with the original food (Data Set S3). In other measures, such as total fecundity, drive inheritance, and viability among the progeny of male drive carriers, no significant differences were found compared to the first batch of crosses. The reduction in the drive inheritance rate among adults suggests that drive-carrying eggs suffered from lower viability than eggs that failed to inherit the drive, indicating a possible viability cost in dry conditions. This could be a direct cost, or it could potentially have occurred by disruption of *yellow-g* in germline cells by drive conversion before they provided sufficient protein for high quality eggs. The latter would potentially explain why there was no noticeable reduction in viability in earlier experiments with more moist food or in the progeny of male drive carriers.

### Cage study

To assess the ability of our homing suppression drive to spread over the course of several generations, we conducted a cage study with a large population size averaging 4,000 individuals per generation (Figure S3). Flies heterozygous for the drive and homozygous for Cas9 were introduced into two cages at frequencies of 41% and 8.8% and were allowed to lay eggs in food bottles inside the cage for one day. Flies homozygous for the Cas9 allele were similarly allowed to lay eggs in separate bottles. We then removed the flies and placed the bottles together in each population cage. The cages were followed for several generations, with all individuals in each discrete generation phenotyped for DsRed to measure the drive carrier frequency, which includes drive homozygotes and heterozygotes. In both cages, the drive carrier frequency increased to approximately 63% (Figure 4, Data Set S4). This possibly represents an equilibrium frequency, though the cages may have increased to a somewhat higher equilibrium frequency with additional generations (models of this drive type predict an asymptotic approach to the equilibrium frequency^24^, making it difficult to estimate from limited cage data). Such an equilibrium result is expected for suppression drives with imperfect drive conversion efficiency. Only when this equilibrium level is high enough will the population actually be eliminated.

**Figure 4.**
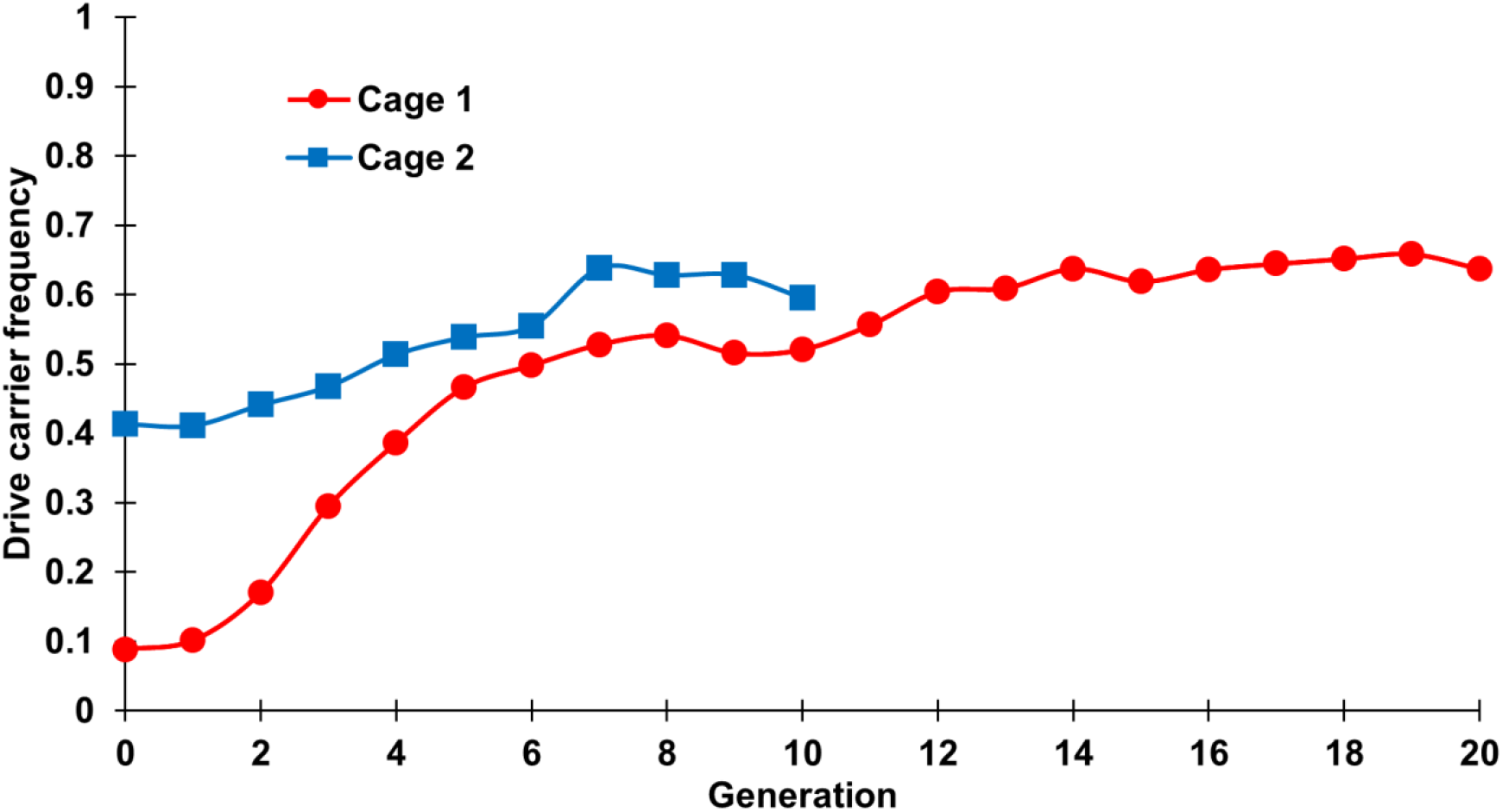
Frequency of drive carriers in cage. Flies carrying one copy of the drive allele and two copies of Cas9 were introduced at initial frequencies of 8.8% (cage 1) and 41.3% (cage 2) into a population that was wild-type at the drive site and homozygous for the Cas9 allele. The cage populations were followed for several non-overlapping generations, each lasting twelve days, including one day of egg-laying. All individuals from each generation were phenotyped for DsRed, with positive drive carriers having either one or two drive alleles (all drive carriers in the initial generation were drive/wild-type heterozygotes).

However, given the average drive conversion rate in heterozygotes of 76.7%, the equilibrium value seen in our cages is substantially lower than the expected drive carrier equilibrium frequency of approximately 90% for a simple model of homing suppression drives with one gRNA^24,44,47^ (a more advanced model with different drive conversion rates between sexes and multiple gRNAs would predict a marginally higher equilibrium frequency^24^). In these models, such a reduction in equilibrium frequency could be explained by a fitness cost of approximately 20% in drive homozygotes (with multiplicative fitness costs and assuming the drive would slowly increase to an equilibrium carrier frequency of 70%)^24^. While the cage experiment food started out moist as in our first set of individual crosses, it was exposed to the air throughout the experiments, potentially resulting in similar or even higher fitness costs to female drive egg viability than those seen in the individual crosses with drier food. Nevertheless, the drive frequency did not decrease systematically over the course of the experiment after the initial increase of the drive, suggesting that functional resistance alleles did not form at a high rate. This is in contrast to four recent studies (three of homing suppression drives^26,30,37^ and another of a modification drive with a costly target site^33^) where functional resistance alleles outcompeted the drive alleles despite higher drive efficiency and lower overall resistance allele formation rates.

A substantial reduction in the population size for this homing suppression drive was not observed (Figure S3). This is likely due to the modest genetic load of the drive, which we use as a measure of the reduction of reproductive capacity of a population (0 = no loss of reproductive capacity compared to wild-type, 1 = the population can no longer reproduce). The load of our drive is closely related to the proportion of sterile females in the population, which increases with drive frequency. In our cages, drive frequency only reached a moderate level that was likely fairly close to its equilibrium level, thus imposing only a moderate genetic load. This genetic load was lower than expected because the drive appeared to carry a fitness cost of unknown type (aside from the intended fitness effects causing sterility in females lacking a functional copy of *yellow-g*), which would directly reduce the drive’s equilibrium frequency and genetic load^56^. Additionally, the drive would likely require a particularly high genetic load to reduce the population at all due to the robustness of the cage population^57,58^. Specifically, the flies likely laid an average of over 20 eggs per female (Data Sets S1-S3), and reduced larval densities usually leads to healthier adults (which could perhaps mature faster and lay even more eggs due to greater size obtained as larva). Indeed, with the low competition found in our vials (Data Sets S1-S3), egg to adult survival was approximately 80%, thus potential enabling a cage population to remain high if even just one out of eight females remain fertile. Reducing female fertility by this amount would require a genetic load of 0.875, and the predicted genetic load in our drives in the last several generations of our cages was perhaps slightly higher than 0.5^24^. While the drive may still have caused a small population reduction in our cages, this was not detectable given the level of fluctuation in population sizes between generations, which could have been caused by variation in larvae density and other random factors.

To further investigate the nature of the drive’s fitness costs, a new cage was established with flies both homozygous and heterozygous for the drive allele but lacking the Cas9 allele required for homing. Such flies were placed in a cage at an initial drive carrier frequency of 76%. Over ten generations, the drive-carrier frequency declined to 29% (Figure S4). This observation is consistent with the drive allele being a recessive female sterile allele (as expected from its disruption of *yellow-g*) and having no additional fitness costs beyond female homozygote sterility in the absence of a genomic source of Cas9. The previous experiments indicated low somatic expression and similar fecundity of drive flies compared to *w*^*1118*^ flies. Thus, all or most of the unknown fitness cost apparently requires the drive allele to be combined with Cas9.

### Maximum likelihood analysis of fitness from cage data

To computationally assess drive performance, we adapted a previously developed method^25,52^ to infer fitness costs based on phenotype data from population cages. We used a simplified model that included only a single gRNA and initially neglected possible formation of functional resistance alleles, assuming that all resistance alleles were nonfunctional (Figure S5). Note that this simplifying assumption of one gRNA for a drive with four slightly underestimates drive performance compared to a more complex model^24^ with the same parameters for drive-wild-type heterozygotes. Since drive carrier individuals in the initial generation of all three cages apparently had substantially lower fitness than in other generations (most likely due differences in health in the populations of the initial generation, though assortative mating could also partially explain the observed effect), likelihood values for the transition from the initial generation were excluded from the analysis.

We reason that the first cage is more reliable for parameter estimation due to the greater number of generations and lower starting frequency, allowing more generations in which the drive can increase toward its possible equilibrium frequency. This equilibrium is predicted by models of homing suppression drives that match the design of our drive^24,44,47^. In this cage, a model of viability-based fitness costs had the best fit to the data based on the Akaike information criterion corrected for small sample size^52^, with drive homozygotes having a viability of 80% (95% confidence interval: 72-88%) compared to wild-type individuals (Table S4A). We did not observe this reduced viability in our assays based on individual crosses (which only showed a modest reduction in viability in offspring of females with the drive when food was dry), but these had limited power to detect such reduction in drive heterozygotes. More importantly, individually assayed flies probably did not experience the same intense competition that might be found in the cage populations, which also were open to the air and perhaps even drier than our second set of individual cross experiments, so modest fitness effects in these cages is quite plausible. A model that included reduction of female fecundity and male mating success matched the data nearly as well as the viability model.

In the second cage, a model with fitness costs from somatic Cas9 cleavage of *yellow-g* in female drive/wild-type heterozygotes was the best match to the data (Table S4B), with such females having a 57% reduction in fecundity. However, this result is not consistent with our direct measurements of fecundity for drive heterozygous females, which could likely have detected such a large fecundity reduction (Data Sets S1-S3). A model with off-target viability fitness costs due to Cas9 cleavage of distant sites was the next best match, though this model did not perform well in the first cage.

Combining data from the two cages (Table S4C), the best model remained one based on somatic Cas9 cleavage and fitness costs. However, models with direct viability and fecundity/mating costs were nearly as good of a match to the data. In a control cage lacking Cas9, the drive declined as expected for an allele that caused recessive sterility in females, with no additional fitness costs (Table S4D). This indicates that any fitness costs are likely mostly due to the drive itself together with Cas9, rather than strong haploinsufficiency of *yellow-g* or an effect in males. Overall, while our analysis provides strong evidence for a fitness cost compared to the neutral model (in agreement with our finding of a lower than expected equilibrium frequency), we were unable to determine the exact nature of the cost. Despite our high sample sizes in each generation, this result is within expectations based on our previous exploration of fitness cost inference in maximum likelihood models, given the autosomal genomic loci and lack of particularly strong fitness effects in this drive system^52^.

### Assessment of functional resistance allele formation

We did not observe qualitative evidence of functional alleles in our cages, which would have resulted in a systematic decline in drive frequency toward the end of the experiments when present at sufficient frequency. Furthermore, our maximum likelihood method allows us to estimate an upper bound of the maximum rate at which functional resistance alleles may have been formed in our population cages. To accomplish this, we allowed the “relative r1 rate” to vary in addition to fitness cost for the two best fitness models, assuming the functional resistance allele formation rate to be proportional to the nonfunctional resistance allele formation rate. The most likely estimate for functional resistance allele formation was 0%, so adding the new r1 parameter produced a substantially worse match by Akaike information criterion (Table S4E). The two cages together had an 95% confidence upper bound of 0.3% for the relative r1 formation rate (fraction of total resistance alleles that were functional), which corresponds to germline and embryo functional resistance allele formation rates of 0.067% and 0.16%, respectively. Note that computational models predict even lower rates of functional resistance allele formation in 4-gRNA drives, likely below 0.01% the level of the nonfunctional resistance allele formation rate, perhaps even lower by several orders of magnitude if the rate of functional sequence repair at individual sites is well below 10%^24^.

Because functional resistance alleles are expected to be rare, normal cage studies have limited power to detect them if they remain at low frequencies. In our population cages, the highest genetic load was about 0.5, meaning that functional resistance alleles at low frequencies would perhaps have a 2-fold advantage over other alleles that would be at equilibrium, preventing them from substantially influencing drive carrier trajectories for the first few generations after they form and thus limiting power to detect them without continuing the cage for several generations. To assess functional resistance alleles for single-sex fertility systems with low drive conversion efficiency, we modified a protocol developed previously for detecting functional resistance alleles in cage studies^30^ by artificially increasing genetic load (see methods for details). Because non-drive carrying flies are removed in this method for each generation, the equilibrium genetic load on the populations was close to 98% according to our previous model^24^ based on the same parameters we used in maximum likelihood analysis of the cage. Functional resistance alleles would retain high fitness if they form, even if those not inherited with a drive allele are removed. Equilibrium genetic load is expected to be reached on the second generation. On the first generation afterward, the predicted genetic load would still be 96%. If any functional resistance alleles are present, they would quickly reach high frequencies, resulting in many flies quickly becoming drive or functional resistance allele heterozygotes and drive homozygotes, avoiding population suppression due to high genetic load. Sequencing a few of these flies would allow for characterization of the functional resistance alleles.

However, while population sizes in the first generation were large in three replicates (Table S5) (245, 250, and 271, of which a handful did not carry a drive allele), the population sizes in the next generation were smaller (84, 93, 43). In the following generation, the population was 21 and 16 in the first two groups and zero in the third group. There were no offspring from the remaining two replicates. This indicates that no functional resistance alleles formed during the study or were quickly lost without contributing to the next generation. Modeling indicates that a relative r1 rate (fraction of total resistance alleles that preserve the function of the target gene) of 0.1% would have resulted in formation of an functional resistance allele by the second generation in about half of cages^24^, likely saving the population from suppression. Thus, the actual relative r1 rate is most likely below 0.1%, consistent with our maximum likelihood results.

## DISCUSSION

This study experimentally demonstrated the utility of gRNA multiplexing as a means for improving the ability of a homing suppression drive to spread through a population without significant formation of functional resistance alleles. The drive displayed a higher drive conversion rate than most single gRNA *Drosophila* drive systems^19,20,22^, as well as a previous homing suppression drive with four gRNAs^11^. However, it had a moderate embryo resistance rate that presumably reduced its rate of spread through the cage. Furthermore, our analyses suggest that the drive also carried a fitness cost of unknown origin. Because drive conversion efficiency was imperfect, these factors together reduced the genetic load of the drive on the population, ultimately preventing elimination of the large cage populations. Indeed, our cages were the first in which a suppression drive failed due to inadequate genetic load rather than functional resistance alleles, underscoring the need for suppression drives to be highly efficient.

Notwithstanding, this study still demonstrated an additional strategy against functional resistance allele formation in suppression drives that is complementary to the targeting of highly conserved sites, which was previously demonstrated as an effective approach^32^. Indeed, all other drives targeting essential or highly important genes without rescue suffered from functional resistance allele formation^26,30,33,37^, which our drive avoided, despite the large population size in our cages. Combined, these two strategies for reducing functional resistance alleles would likely be even more effective while still maintaining high drive conversion efficiency^24^.

Since genetic load (which determines the suppressive power of a drive) is mostly determined by drive conversion efficiency and fitness costs, this represents a hurdle for any suppression strategy based on a homing drive. As the frequency of the drive increases, so does the rate of drive removal. With 100% homing efficiency, the relative frequency of the drive allele would continue to increase as the population numbers decline, and complete suppression would occur as the drive reaches fixation. However, with a lower efficiency, wild-type alleles remain, and the antagonistically acting forces of drive conversion and drive allele removal result in an equilibrium frequency. Fitness costs from the drive would, in this case, further reduce the equilibrium drive frequency and resulting genetic load. Homing drives in *Anopheles*^27,32,37^, even with similar design and promoters, have demonstrated consistently higher drive conversion rates compared to drives in fruit flies. In mosquito suppression drives with the *zpg* promoter, the higher somatic fitness costs were more than compensated for by a higher drive conversion efficiency, resulting in a superior genetic load. Engineering sufficiently high drive conversion efficiency could therefore be a challenge when designing drives designed to be employed for suppression in the fields of Drosophilids such as the fruit pest *Drosophila suzukii*.

Additionally, the reduced equilibrium frequency in flies compared to *Anopheles* mosquitoes represents a limitation in our study for detecting functional resistance alleles. At a lower equilibrium frequency, functional resistance alleles have a reduced fitness advantage compared to drive and wild-type alleles, reducing our power to distinguish them using our maximum likelihood method that analyzes drive carrier frequency trajectories. Our analysis of functional resistance alleles also somewhat depended on the fit of our model with drive efficiency (measured from individual crosses, though the actual efficiency may be slightly higher due to the multiplexed gRNA design) and fitness parameters, the latter of which is inferred by the model and may be particularly difficult to accurately assess. Nevertheless, our results showcase the utility of the maximum likelihood analysis for predicting functional resistance based on cage phenotype frequency trajectories without additional sequencing. However, such sequencing of each cage generation would still likely have allowed for greater power in detecting functional resistance alleles. Our use of the artificial selection small cage study with a modified protocol represents a potential way to detect functional resistance alleles more directly, with far higher power given an identical experimental effort invested. This method was based off a previous *Anopheles* study^30^, but modified to retain high power to detect functional resistance even when drive conversion is lower and when the phenotype involves single-sex sterility instead of both-sex reduced viability.

It remains unclear exactly what caused the fitness costs associated with our homing suppression drive. Though we could estimate the magnitude of such costs, there is considerable uncertainly in their exact value. There may be additional factors at play that we did not model due to lack of evidence in this or other studies, such as potentially reduced inheritance of cleaved chromosomes that would have the tendency of inflating calculated drive conversion, which would reduce apparent fitness costs compared to actual costs. The nature of the fitness costs also could not be determined based on our data. It is possible that a combination of several different types of fitness costs was at play, including fitness components that we did not include in our model. For example, perhaps the target gene was slightly haploinsufficient, causing females with only one wild-type allele to have slightly reduced fertility compared to wild-type females. Another possibility is that *yellow-g* is partly required by germline cells that were underdoing drive conversion (thus eliminating their remaining wild-type allele), which could explain the reduced egg viability seen in some of our individual crosses with drier food (where reduced levels of *yellow-g* below that provided by a single, stable allele could weaken the egg casing, making the eggs more vulnerable to dry conditions). These issues could potentially be addressed by changing the target gene to one of many other possible female fertility genes that does not have a germline-related maternal effect. Another possibility is off-target cleavage effects as seen previously^53^, which would likely be exacerbated by multiplexed gRNAs. However, such an issue could be addressed relatively easily by using high fidelity Cas9 nucleases that show little to no off-target cleavage^59–65^, which have been shown to have similar drive performance^53^. By contrast, if fitness costs are caused by the expression of the drive components themselves, they may be more difficult to directly address. In this case, increasing drive conversion efficiency, for example by using a different Cas9 promoter, may be the best route to developing successful drives, while potentially also minimizing fitness costs from direct component expression by further limiting expression only to cells and time windows where drive conversion takes place. Indeed, modeling indicates that high drive efficiency and fitness may play an even more important role in ensuring success in complex natural populations with spatial structure^45,50,58,66,67^.

If the drive conversion rate cannot be sufficiently increased in a *Drosophila* or other species given the set of available genetic tools, then a non-homing TADE type suppression drive^68,69^ may still be able to provide a high genetic load. This only requires high efficiency for germline cleavage (regardless of whether it results in homology-directed repair or end-joining) rather than for the drive conversion process (which requires homology-directed repair). Though engineering such a drive targeting haplolethal genes may be challenging, working with such genes is possible at least for homing drives^25^. However, TADE drives are frequency-dependent and thus weaker than homing drives, requiring higher release sizes for success^68,69^. In some cases, this feature may be desirable if the drive should be strictly confined to a target population. Another way to achieve confinement that could still involve a homing suppression drive would be to use a tethered system in which the split homing element is linked to a confined modification drive system^70,71^. Such a method would also allow split homing suppression drive elements, similar to the one we tested, to potentially be release candidates if their performance is sufficient.

Overall, we have demonstrated that even avoiding functional resistance alleles is often insufficient to ensure a high enough genetic load to suppress populations, underscoring the need to develop highly efficient drives. We also showed that gRNA multiplexing is a promising technique for reduction of functional resistance alleles in a homing suppression drive while maintaining relatively high drive conversion efficiency. Since multiplexing gRNAs is a fairly straightforward and flexible process using either tRNA, ribozymes, or separate promoters for each gRNA, we believe this approach has the potential to be applied to a wide variety of suppression gene drive designs, providing similar benefits in many species.

## DATA AVAILABILITY

The data underlying this article are available in the article and in its online supplementary material. All supporting code is available on GitHub (https://github.com/MesserLab/HomingSuppressionDrive and https://github.com/MesserLab/Binomial-Analysis).

## ACKNOWLEDGEMENTS

This study was supported by the National Institutes of Health awards R21AI130635 to JC, AGC, and PWM, award F32AI138476 to JC, and award R01GM127418 to PWM.

## SUPPLEMENTARY INFORMATION

### Supplementary Methods

#### Plasmid Construction

##### gRNA-tRNA Array

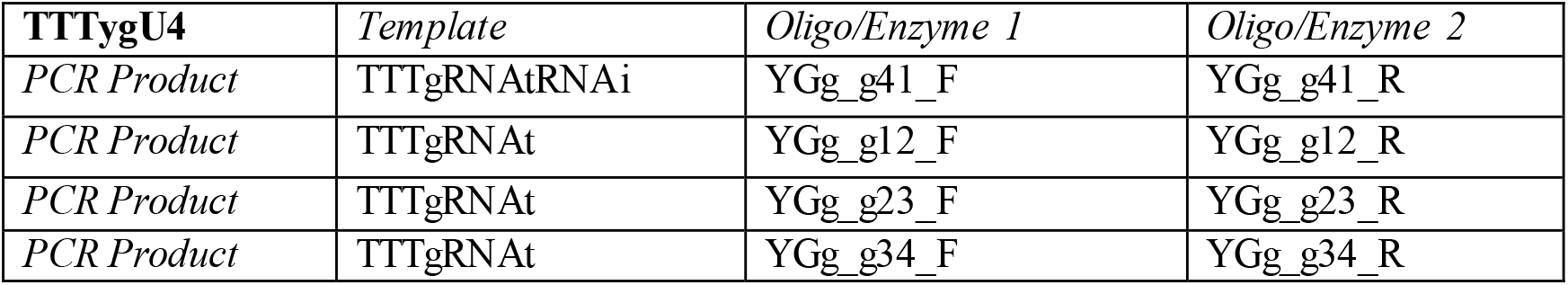

##### Left Homology Arm

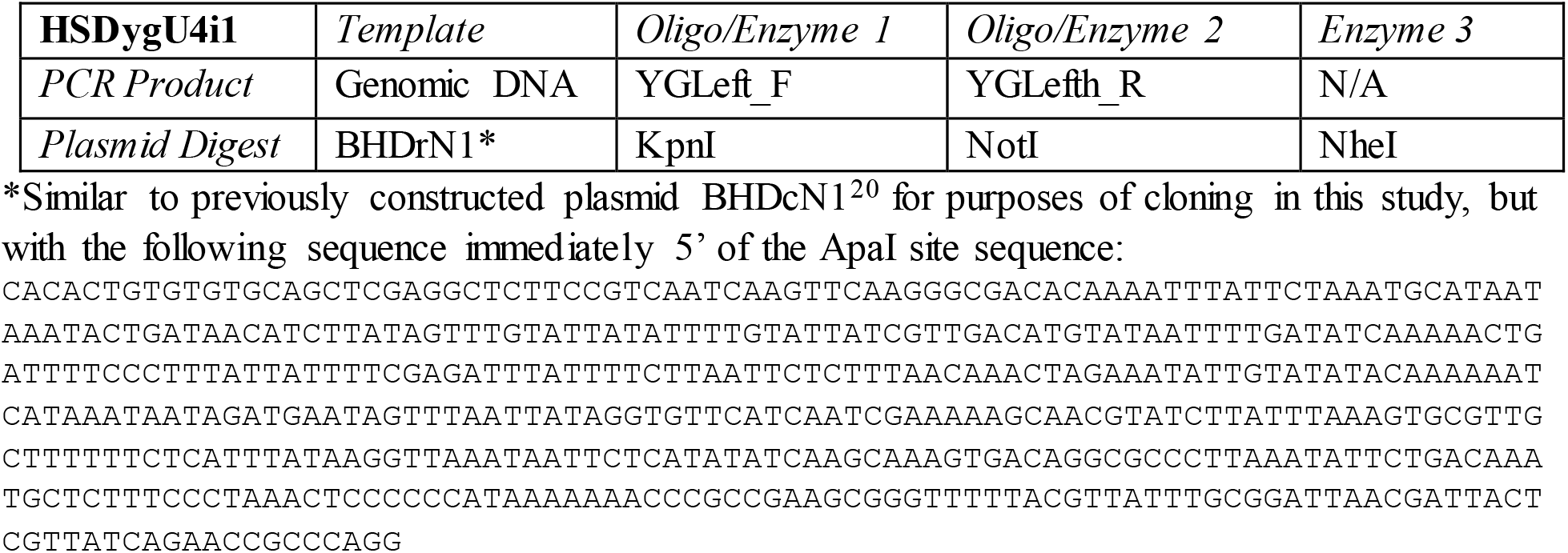

##### Right Homology Arm and gRNA

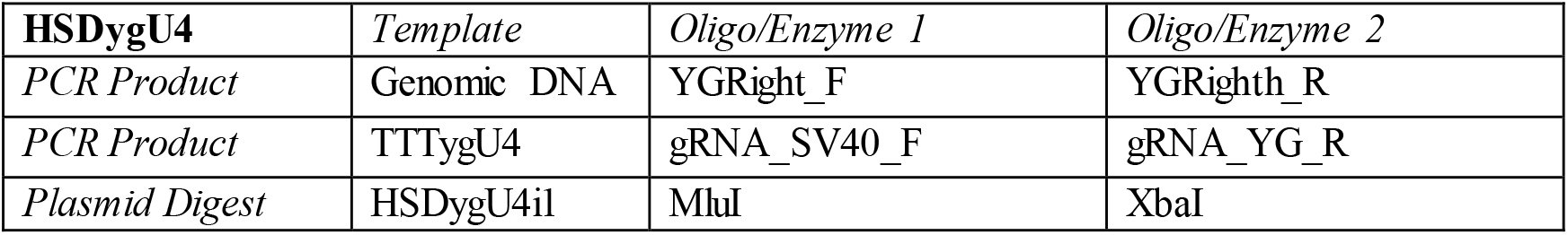

#### Construction primers

##### HSDygU4i1

~~~
YGLeft_F: TAGGGGTCAGTGTTACAACCAATTAACCAGGTACCGTGGGTGGATTACAGGGTAGCA
YGLefth_R: TTAGTCTCTAATTGAATTAGATCCGCGGCCGCCTGCGGATGGGCTGCTCC
~~~

##### HSDygU4

~~~
YGRight_F: TTTAATGTTCGCTTAATGCGTATGCATAGGCCTCCAAGGACAACAAGCCATTCG
YGRighth_R: GGCATCAAACTAAGCAGAAGGCCCCTGACTCTAGAGTGGAGGGATACGGACTCAA
gRNA_SV40_F: GGTTTGTCCAAACTCATCAATGTATCTTAACGCGTTTTTTTGCTCACCTGTGATTGCTC
gRNA_YG_R: CCTATGCATACGCATTAAGCGAACA
~~~

##### TTTygU4

~~~
YGg_g41_F: GTGCACATAAACACGGCCAACCACAGTTTTAGAGCTAGAAATAGCAAGTTAAA
YGg_g41_R: AAAACCAGATGCAGTCCCAAGATCGTGCATCGGCCGGGAATCG
YGg_g12_F: GCACGATCTTGGGACTGCATCTGGTTTTAGAGCTAGAAATAGCAAGTTAAA
YGg_g12_R: AACTGGAGTAGTCGACCACGATGTGCACCAGCCGGGAATCG
YGg_g23_F: GCACATCGTGGTCGACTACTCCAGTTTTAGAGCTAGAAATAGCAAGTTAAA
YGg_g23_R: AACGGTCACCTCCGAGAGTCGGCTGCACCAGCCGGGAATCG
YGg_g34_F: GCAGCCGACTCTCGGAGGTGACCGTTTTAGAGCTAGAAATAGCAAGTTAAA
YGg_g34_R: TGTGGTTGGCCGTGTTTATGTGCACCAGCCGGGAATCG
~~~

#### Sequencing primers (for confirming plasmid sequences and sequencing resistance alleles)

~~~
EGFPaLeft_S_R: GCGAAAGCTAAGCAAATAAACAAGC
U6term_S_F: CATCTGACGTGTGTTTATTTAGAC
Yellow_gRNA1_S_F: TTGCTCACCTGTGATTGCTCC
CFD5_S_R: TAGACAATGGTTTTCCGTTGACGT
YGLeft_S_F: ACAAACGGCAAACAAACGAGG
YGLeft_S_R: TGGCGGCTAATTGAAATGTTGG
YGRight_S_F: TCGAACTGAATCAAGAGTTTGGAG
YGRight_S_R: TGAGCCACACTTCTGAGAACT
~~~

## Supplementary Results

**Figure S1.**
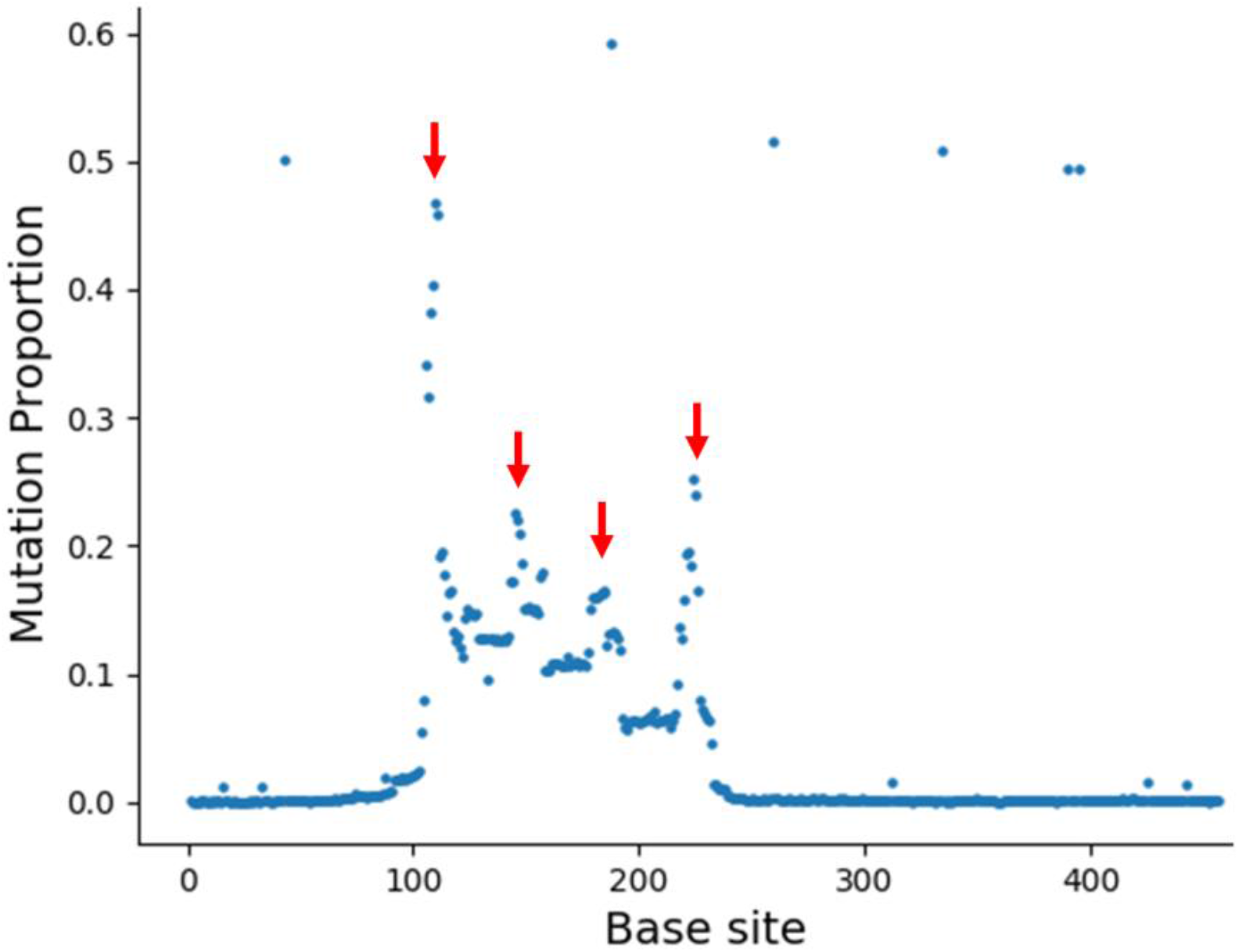
Mutations at gRNA cut sites. Several female and male drive heterozygotes were crossed to each other, and approximately 100 progeny were collected. Pooled DNA was purified and used as a template for PCR around the drive target site. PCR products were analyzed by deep sequencing. The chart shows the fraction of DNA at each nucleotide that was different from the wild-type allele. Arrows show locations of the four gRNA target sites.

**Figure S2.**
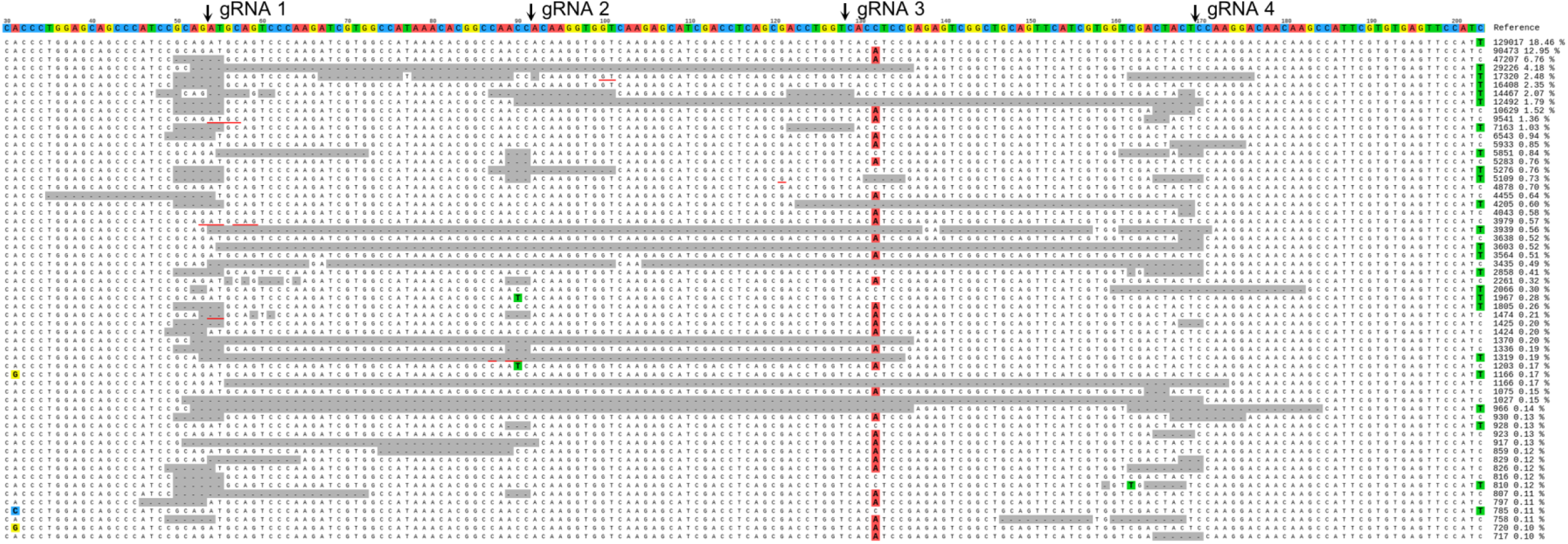
Resistance allele sequences. Several female and male drive heterozygotes were crossed to each other, and approximately 100 progeny were collected. Pooled DNA was purified and used as a template for PCR around the drive target site. PCR products were analyzed by deep sequencing. Arrows above each panel show the gRNA cut sites. Highlighted nucleotides are different from the reference sequence. The “A” variant near gRNA target site 3 may be an error since it was not detected in any Sanger sequencing sample, but it could also represent a pre-existing resistance allele for gRNA #3 present in the population at intermediate frequency. Note that sequences with one or more small inserts interrupting large deletions were likely indels over the entire spanned region with a small insertion that formed during end-joining repair. Thus, the highlighted “T” nucleotides at gRNAs 2 and 3 simply represent sequences of an insertion after end-joining repair.

**Figure S3.**
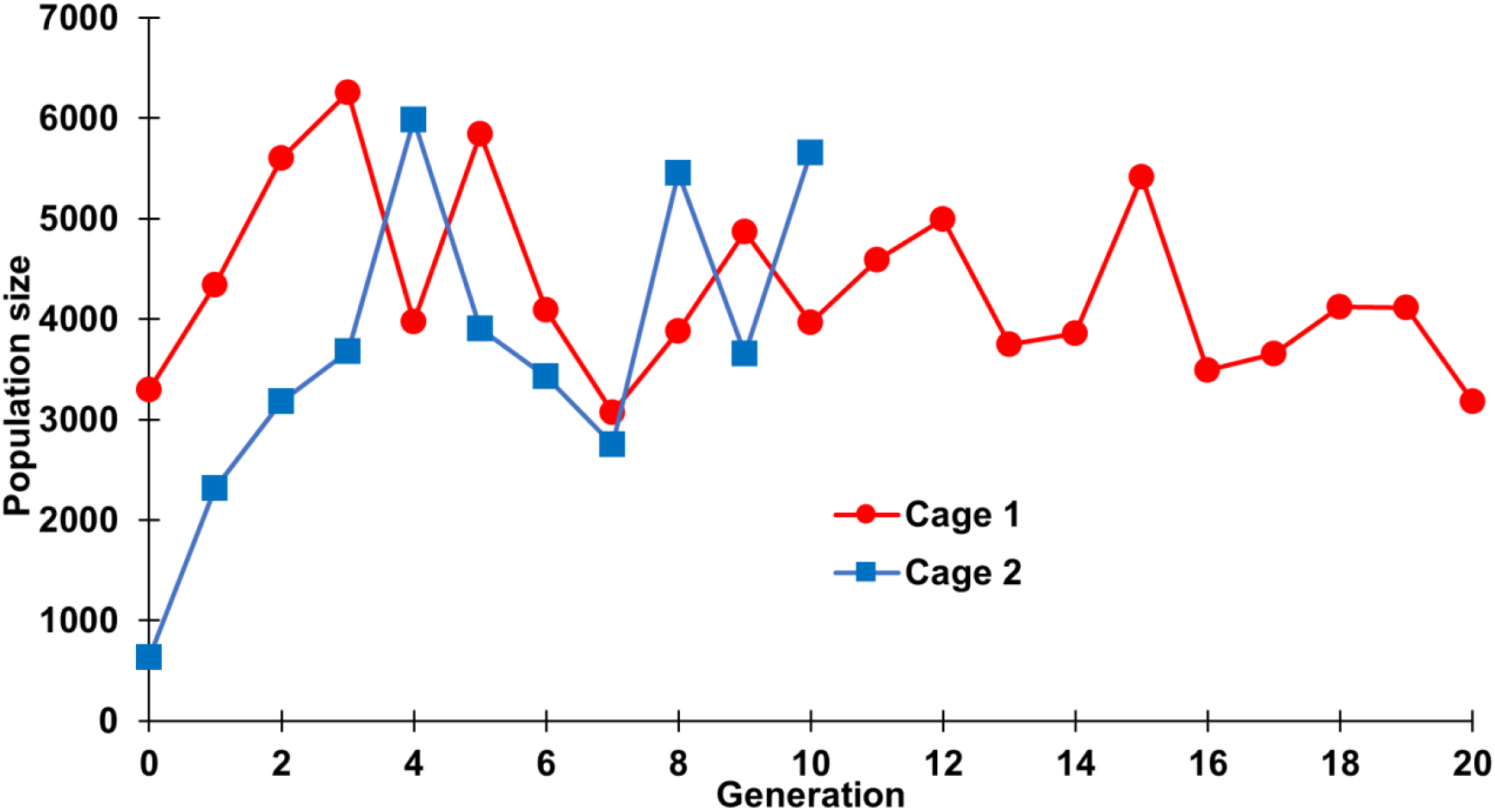
Cage population size. The population size for each generation is displayed for the two drive experiments cages from Figure 4.

**Figure S4.**
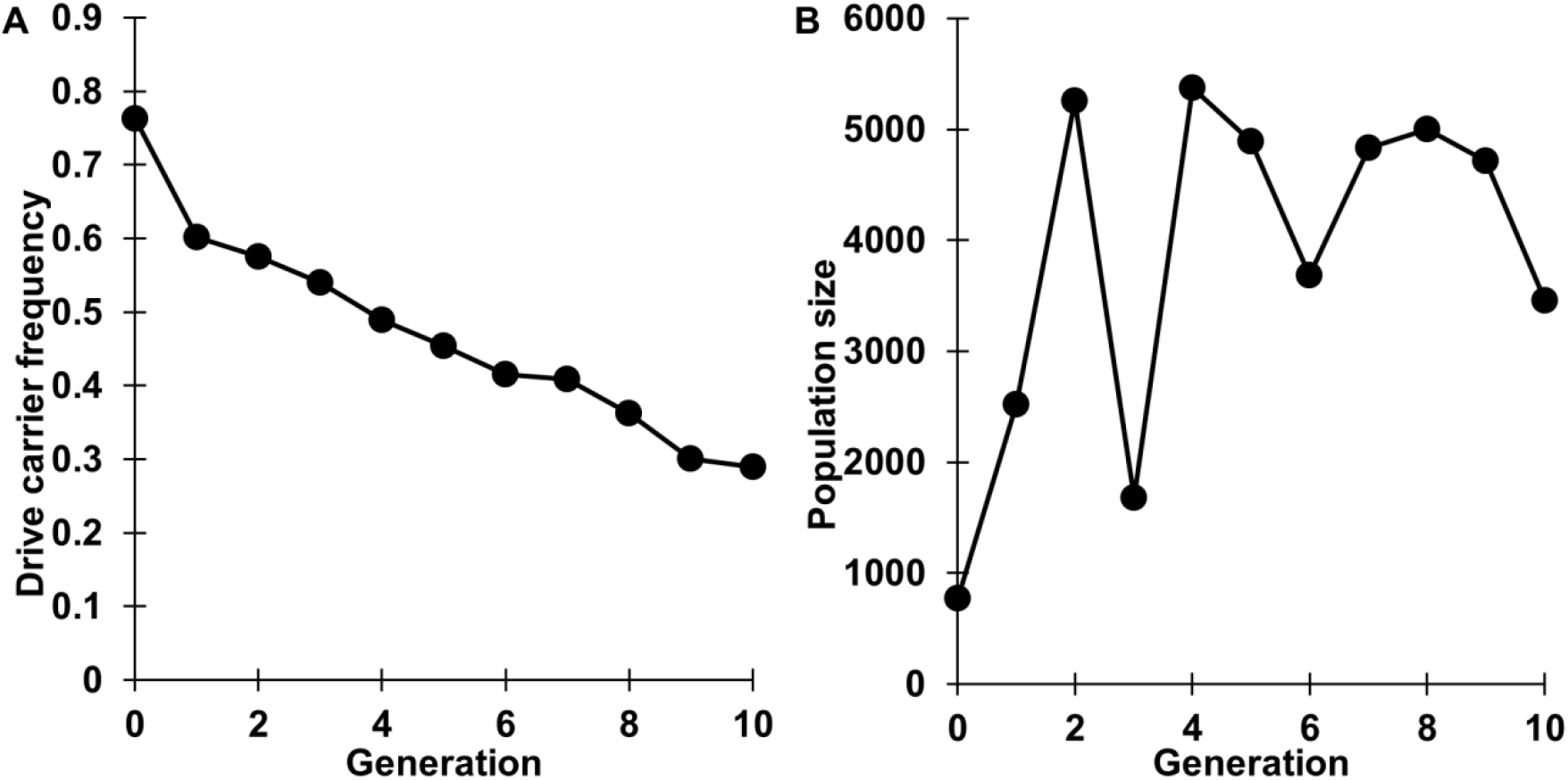
Cage experiment without Cas9. (**A**) Individuals with a drive allele and without Cas9 were introduced into a *w*^*1118*^ cage population without Cas9 at a carrier frequency of 76%. The cage population was followed for several non-overlapping generations, each lasting twelve days, including one day of egg-laying. All individuals from each generation were phenotyped for DsRed, with positive drive carriers having either one or two drive alleles. The drive allele cannot perform drive conversion, so it decreases in the population over time. (**B**) Size of the cage population in each generation.

**Figure S5.**
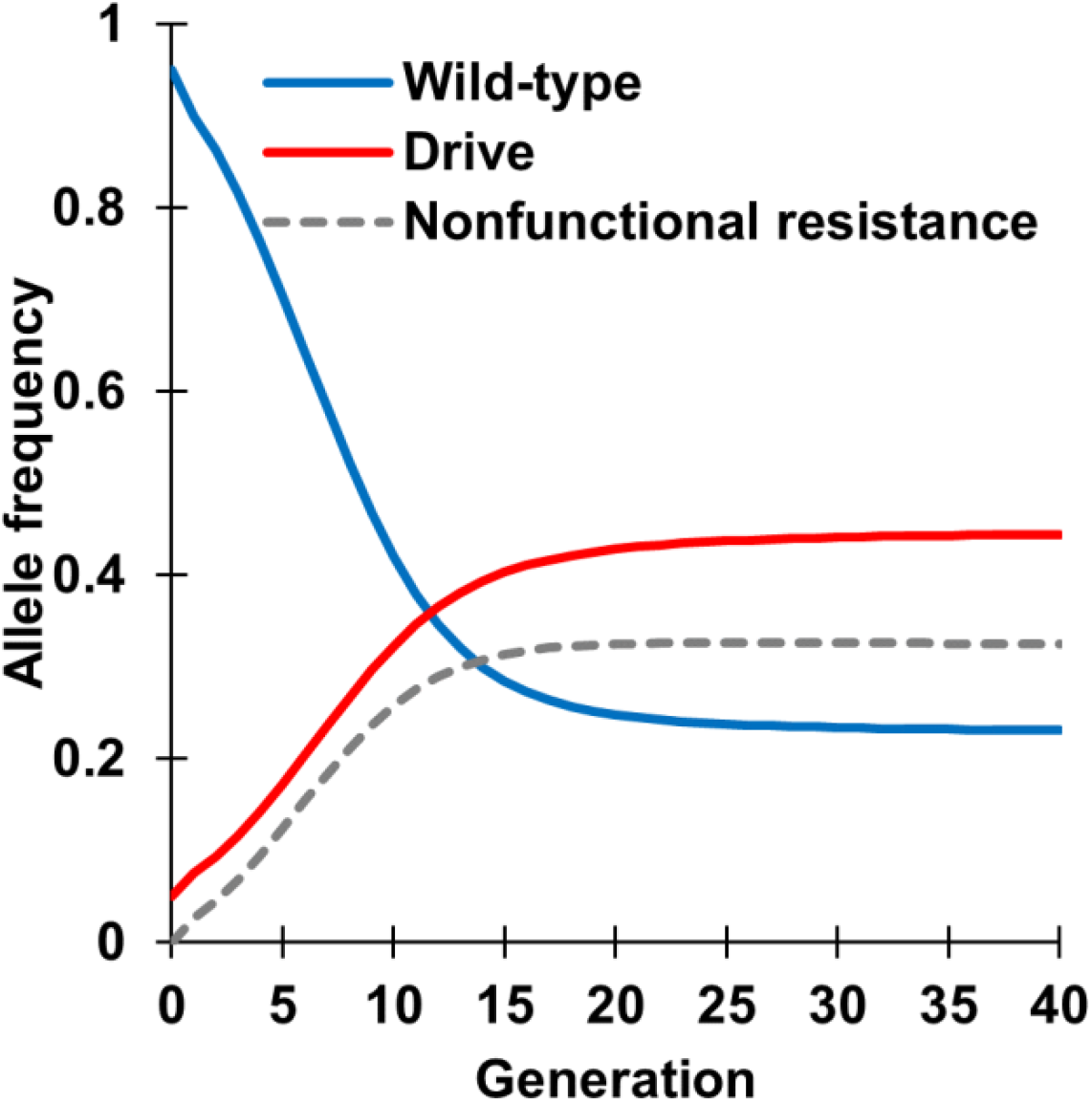
Predicted allele frequencies. Using the same drive efficiency parameters as modeled in our maximum likelihood study and an example female somatic fitness of 0.67, we plotted drive, wild -type, and nonfunctional resistance allele frequency trajectories. Equilibrium is nearly reached after 20 generations.

**Table S1.**
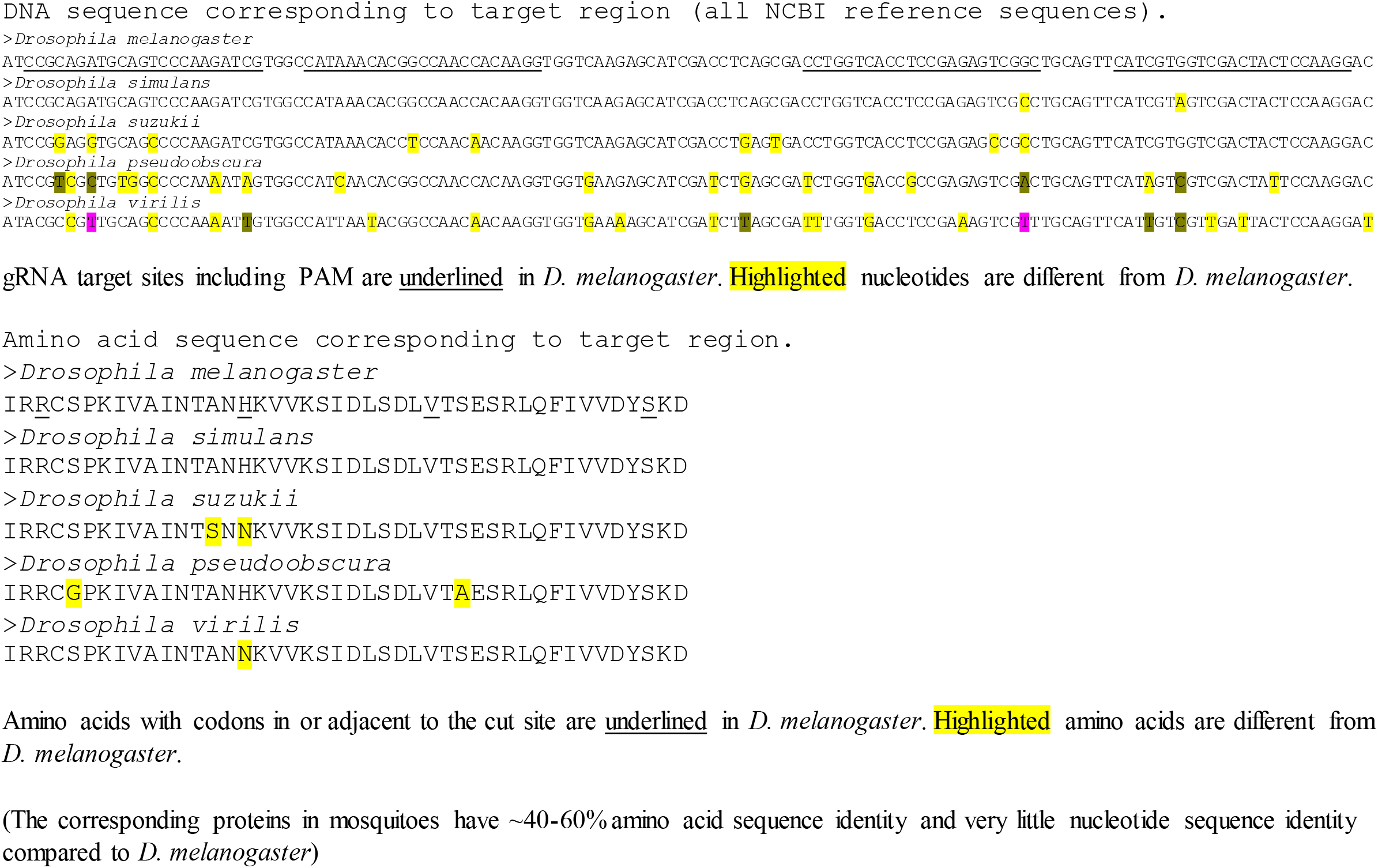
Target site conservation.

**Table S2.**
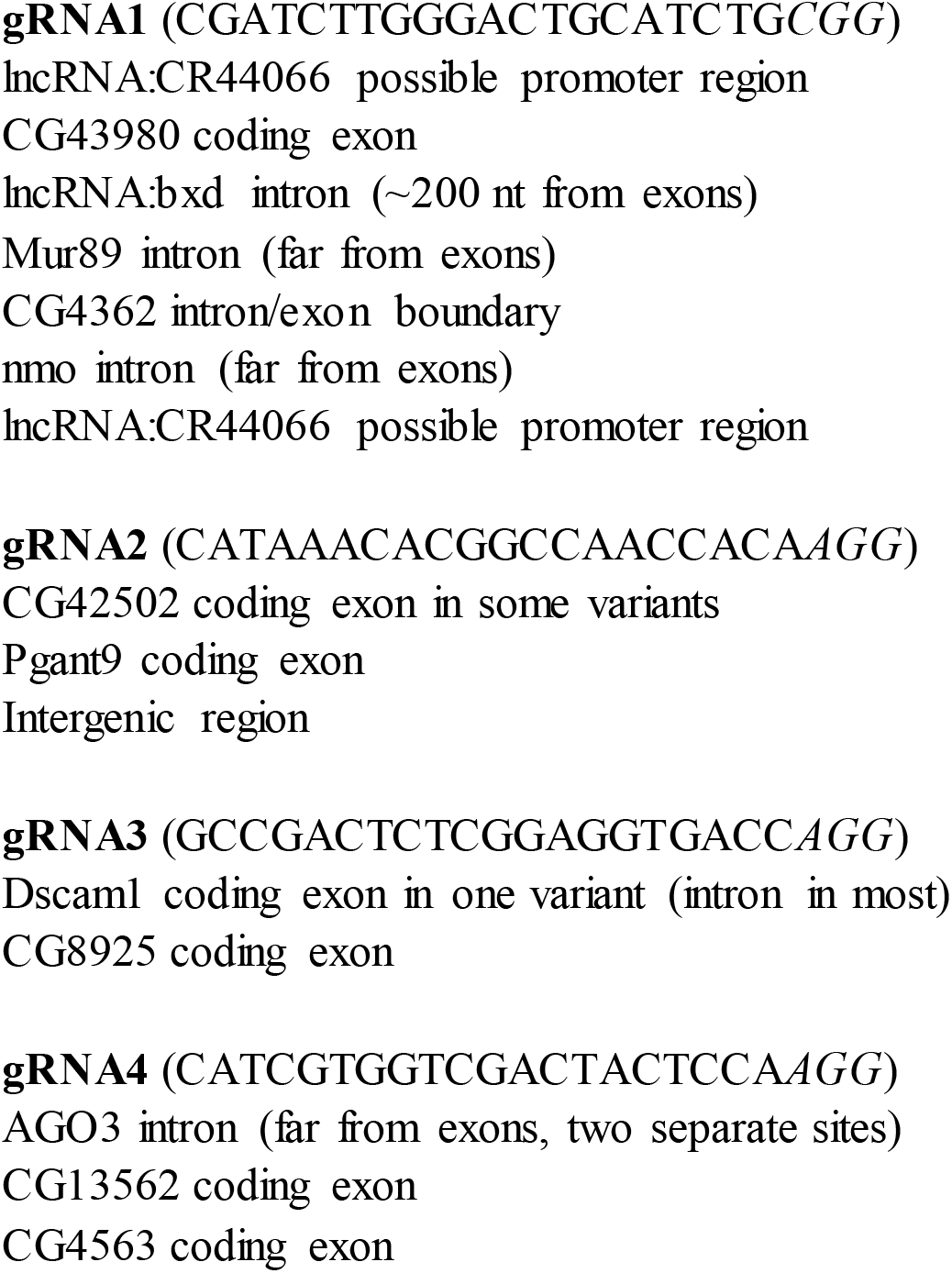
Predicted off-target sites. Off-target sites were predicted with Brown University CRISPR Optimal Target Finder as noted in the drive construct design section. No strong off-target sites were predicted for each gRNA, but several sites were predicted using maximum stringency (gRNAs below are listed in order of their cut site in *yellow-g*).

**Table S3.**
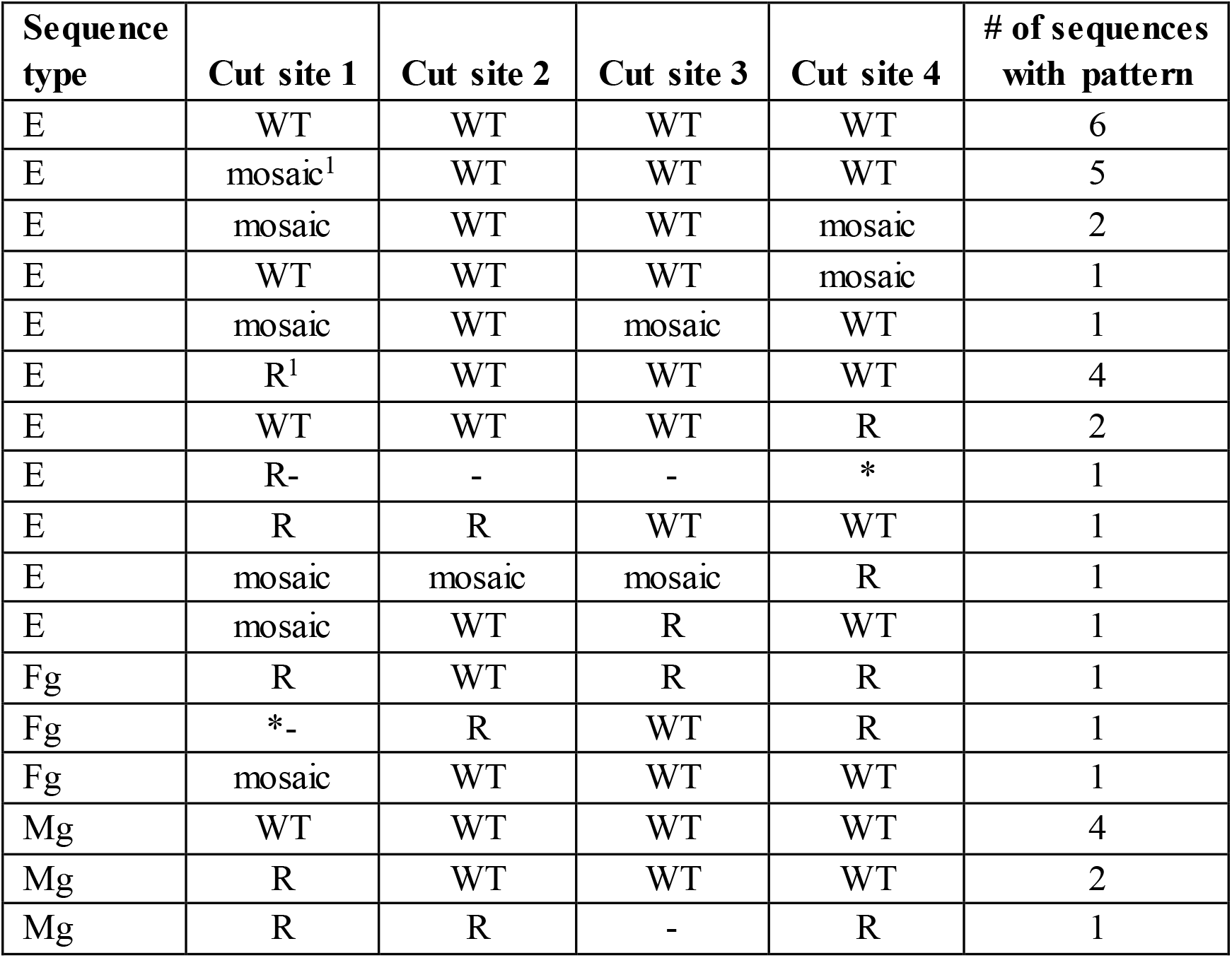
Resistance allele analysis. Progeny of drive heterozygote females or males were sequenced around the target site to identify resistance alleles. For sequence type, “E” indicates that the sequenced progeny is a drive carrier with a drive heterozygote mother. The sequence is therefore an embryo resistance allele. “Fg” indicates that the sequenced individual did not inherit a drive allele from a drive heterozygous mother. These could have two resistance alleles, one from germline and embryo cutting and the other from embryo cutting. “Mg” is similar, but has a male parent, so only germline cutting could take place. For “Fg” and “Mg”, two a **l**eles would be sequenced, and wild-type alleles are not displayed at each cut site unless they are the only one present. WT = wild-type. R = resistance. “-” indicates deletion between the sites. “*” indicates large deletion that continues past the target site. m = mosaic sequence. The high molecular weight product has the sequence displayed above, while the low molecular weight product for both consisted of a large deletion that went well beyond both outer gRNA target sites.

**Table S4.**
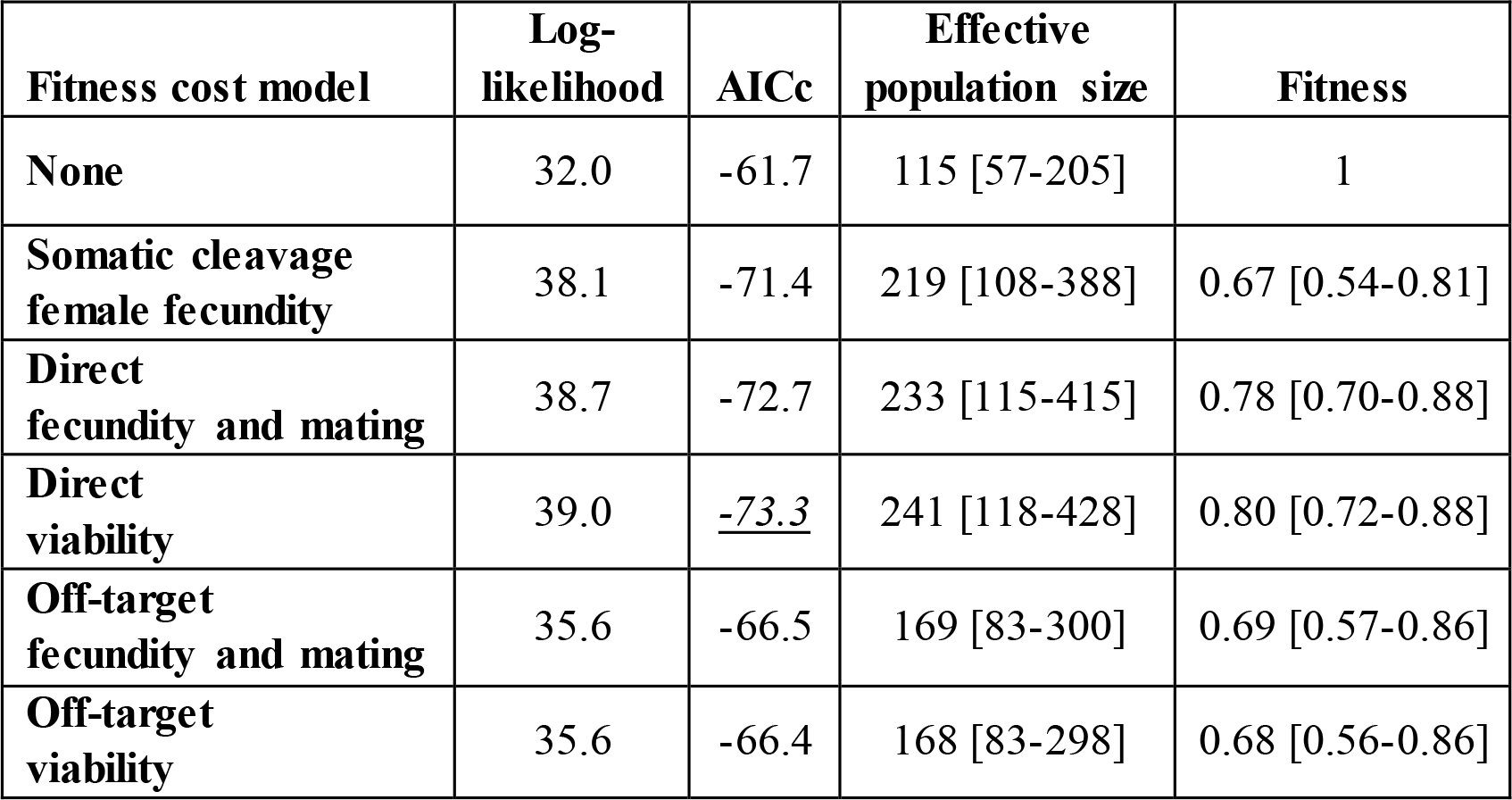
Maximum likelihood parameter estimates from cage populations. Fitness values are for drive homozygotes (multiplicative fitness per allele), except for somatic Cas9 cleavage, where the value is applied directly to drive/wild-type females. 1 is equivalent to wild-type. [Brackets] show 95% confidence intervals. All models have an effective population size parameter, and models with fitness costs have a single additional fitness parameter. Log-likelihood: shows a relative probability (higher values indicate a better model fit) AICc: Akaike information criterion, corrected (low values indicate a better match of the model without overfitting). **Table S4A. Cage 1 drive released into Cas9 background**

**Table S4B.**
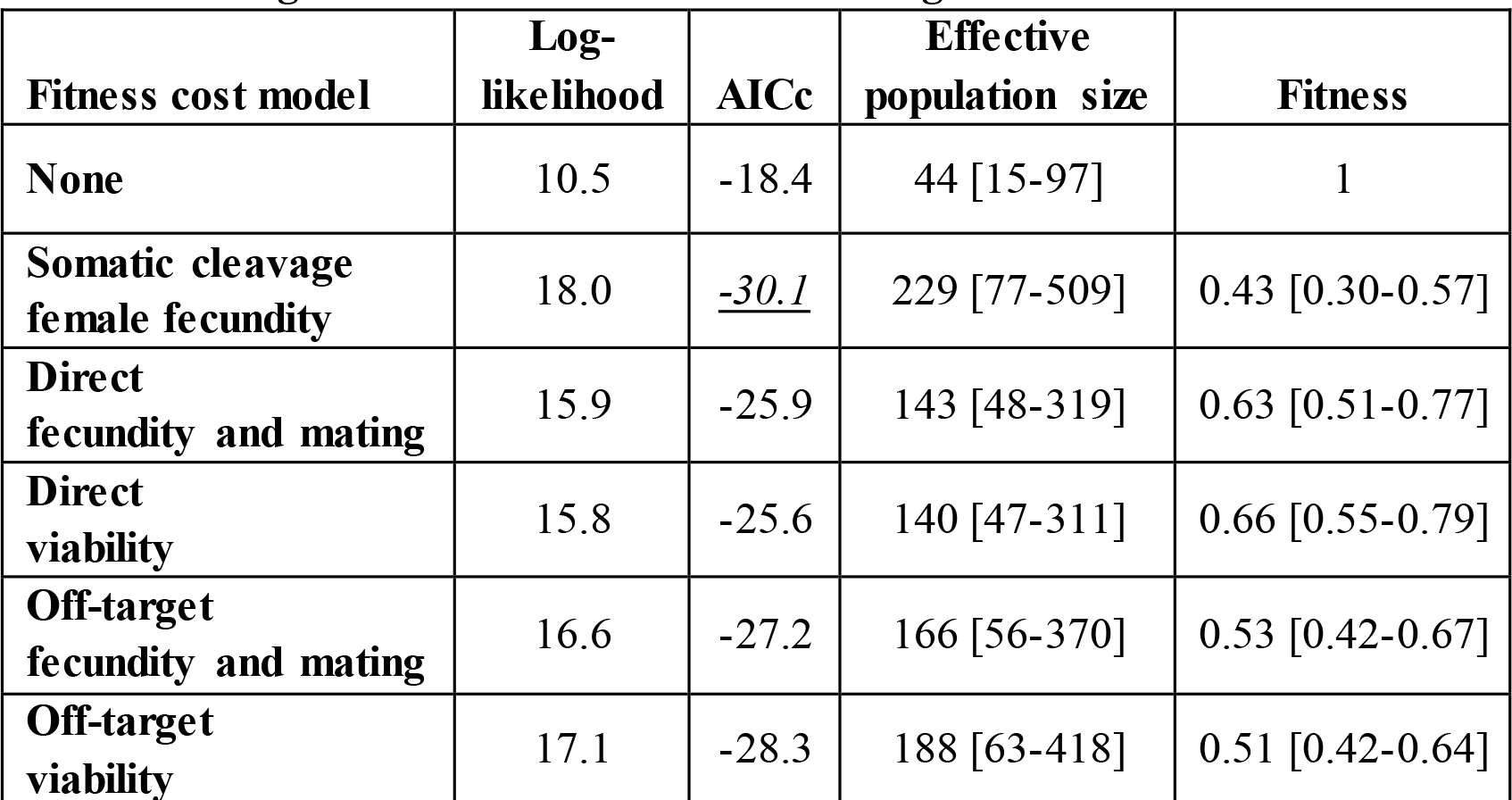
Cage 2 drive released into Cas9 background.

**Table S4C.**
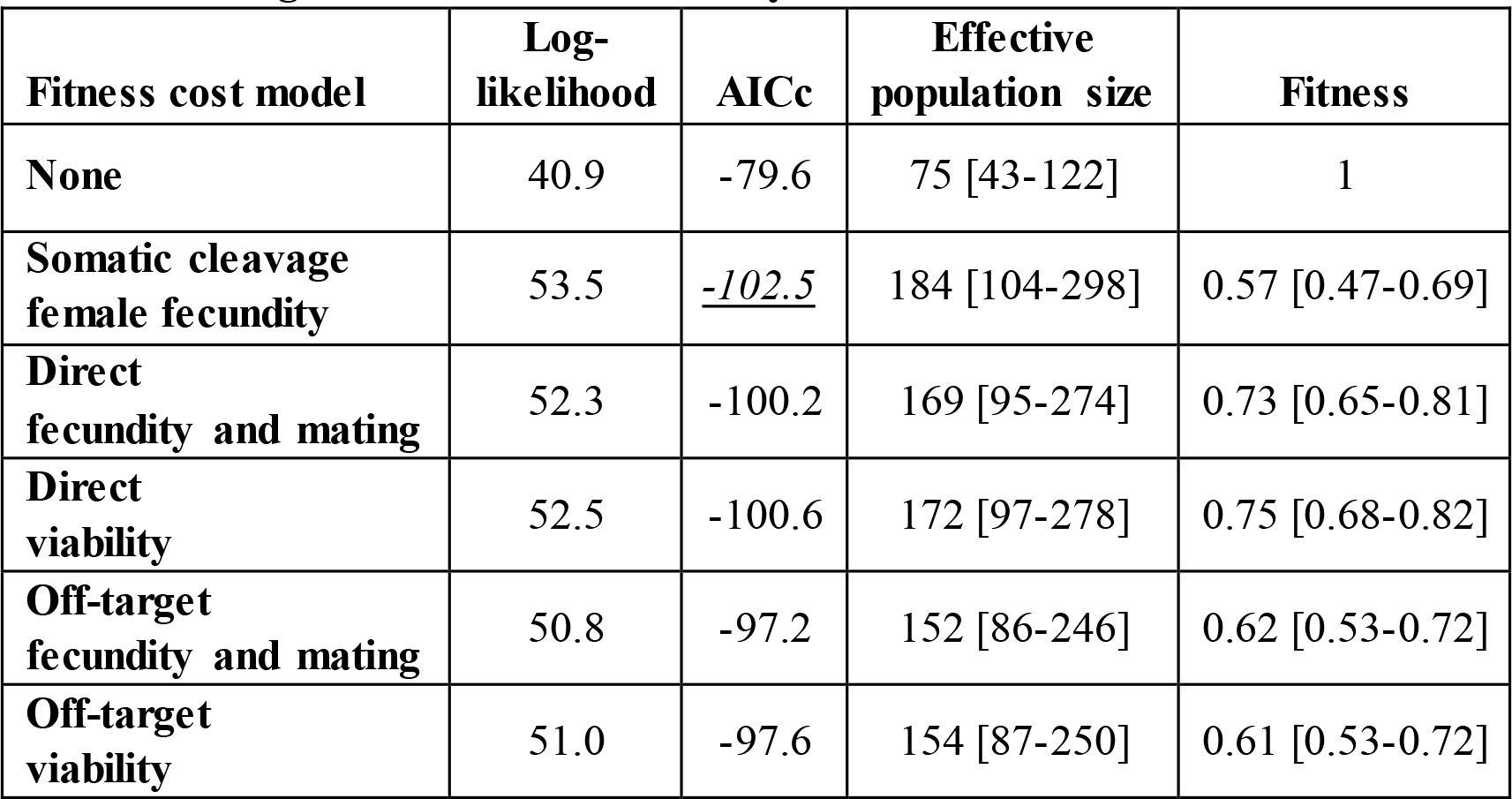
Cages 1 and 2 combined analysis.

**Table S4D.**
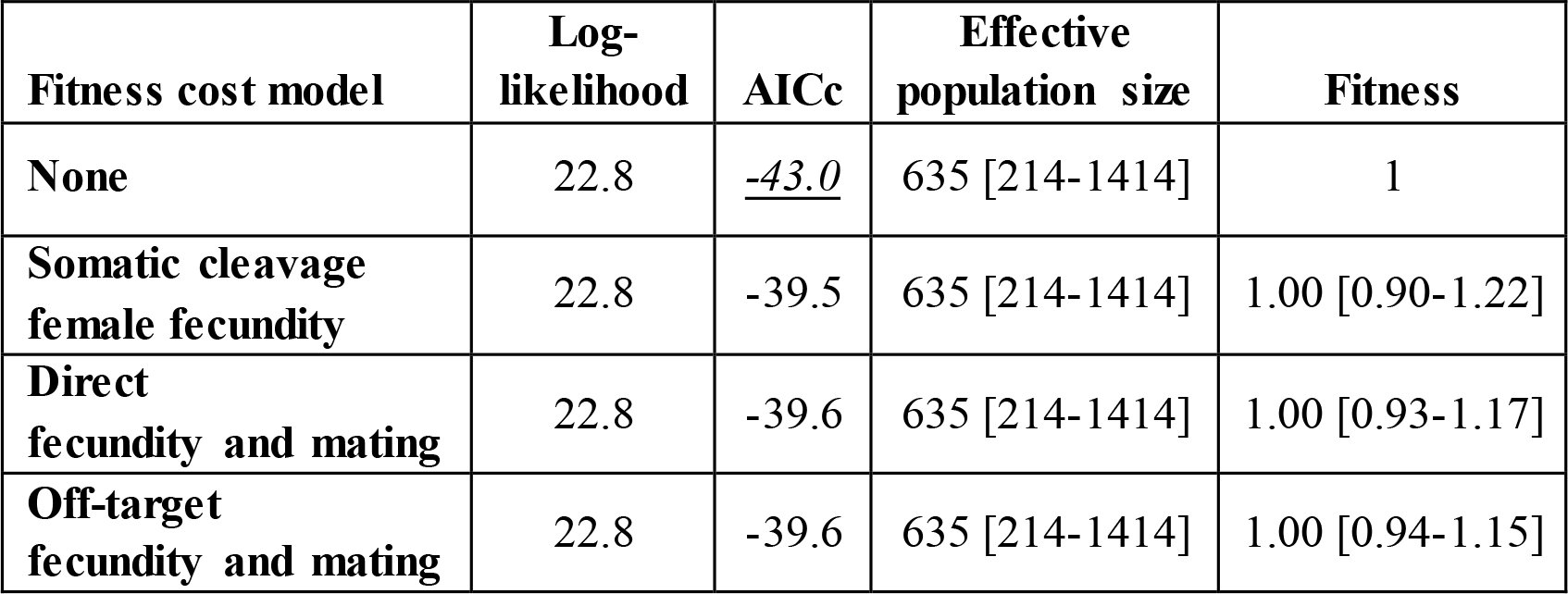
Drive without any Cas9 present.

**Table S4E.**
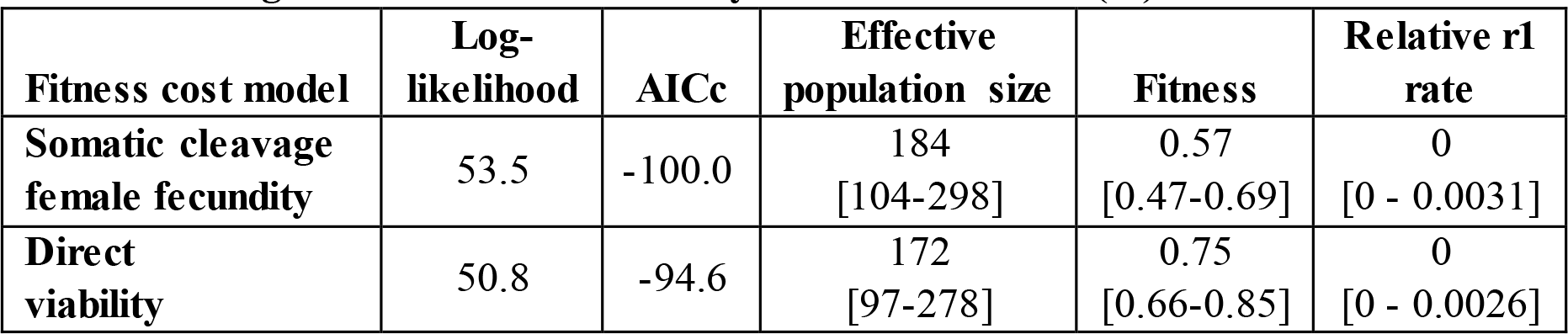
Cages 1 and 2 combined analysis with functional (r1) resistance.

**Table S5.**
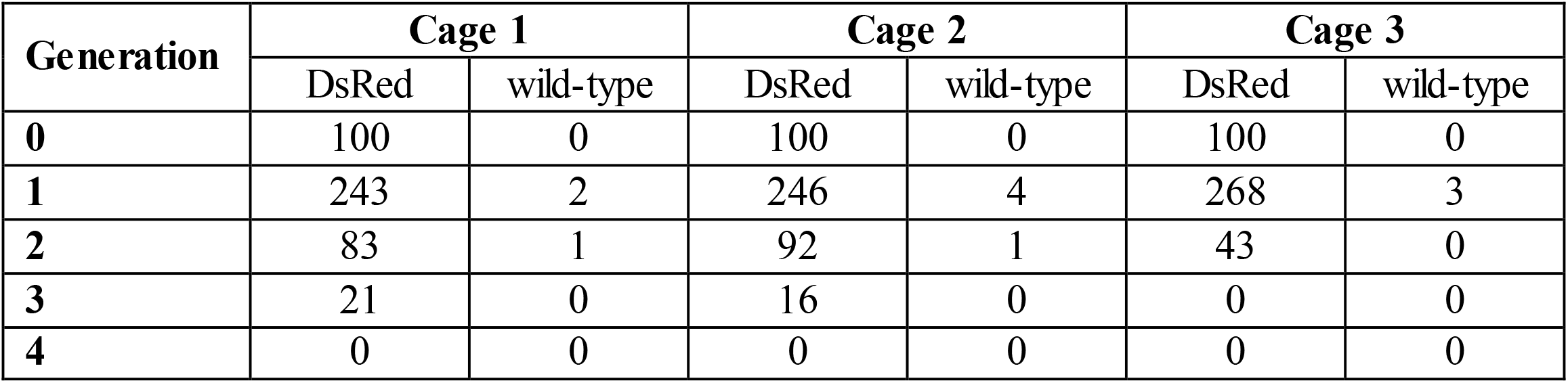
Artificial selection cage experiment.

